# The origin and dynamics of cellular heterogeneity vary across lineage subtypes of castrate resistant prostate cancer

**DOI:** 10.1101/2022.03.24.484651

**Authors:** Michael L. Beshiri, Brian J. Capaldo, Ross Lake, Anson T. Ku, Danielle Burner, Caitlin M. Tice, Crystal Tran, Julianna Kostas, Aian Neil Alilin, JuanJuan Yin, Supreet Agarwal, Samantha A. Morris, Fatima H. Karzai, Tamara L. Lotan, William L. Dahut, Adam G. Sowalsky, Kathleen Kelly

## Abstract

**Purpose:** To resist lineage-dependent therapies such as androgen receptor inhibition in prostate cancer, cancer cells often adopt a stem-like state resulting in lineage-plasticity and phenotypic heterogeneity. We assessed the dynamics of lineage determination and cellular subpopulation expansion in treatment-resistant adenocarcinoma, amphicrine, and small cell neuroendocrine castrate resistant prostate cancers (CRPCs).

**Experimental Design:** We developed CRPC patient-derived organoid models that preserve heterogeneity of the originating tumor, including an amphicrine model harboring epigenetic driver mutations, *ARID1A* and *ARID1B,* and displaying a range of luminal and neuroendocrine phenotypes. We used single-cell RNA-seq, barcode lineage-tracing, single-cell ATAC-seq, and RNA-FISH to delineate the subpopulation structure of the heterogeneous organoids and define the lineage hierarchy, determine potential transcriptional regulators of amphicrine lineage-plasticity, and identify subpopulation-specific molecular targets for therapeutic intervention.

**Results:** Transcriptionally similar stem/progenitor cells were identified for all lineage populations. Lineage tracing in amphicrine CRPC showed that heterogeneity originated from distinct subclones of infrequent stem/progenitor cells that produced mainly quiescent differentiated amphicrine progeny. Amphicrine cells were enriched for secretory luminal, mesenchymal, and enzalutamide treatment persistent signatures. By contrast, adenocarcinoma CRPC had a less defined hierarchy, as progeny originated from stem/progenitor cells and self-renewing differentiated luminal cells. NEPC was composed almost exclusively of self-renewing stem/progenitor cells. Amphicrine stem cells demonstrated concurrent transcription factor activities associated with stem/progenitor, luminal epithelial and mesenchymal lineages. Finally, the amphicrine stem/progenitor subpopulation was specifically depleted with an AURKA inhibitor, which blocked tumor growth.

**Conclusions:** These data illuminate distinct origins and dynamics of subtype-specific CRPC plasticity in addition to demonstrating a strategy for targeting differentiation-competent stem cells.

**Translational Relevance:** For advanced prostate cancer, therapeutic resistance to androgen signaling suppression increasingly involves the development of lineage plasticity. The cellular states of transition and subpopulation heterogeneity that underlie lineage-plasticity are not well understood, which is an ongoing challenge to the design of effective treatments. Using patient-derived organoid models of various CRPC lineage subtypes, we observed distinct patterns with respect to stem/progenitor activity and associated growth phenotypes. The simultaneous expression of AR-driven and neuroendocrine identities, so-called amphicrine tumors, are thought to be an early dedifferentiation stage in plasticity-mediated resistance. We observed in an epigenetically-driven, amphicrine model of CRPC that a rare but necessary bipotent stem/progenitor population is suppressed by AURKA inhibitors, leading to tumor regression, while ARPC demonstrates both self-renewing differentiated luminal cells and stem/progenitors. These data suggest that AURKA inhibition may block the amplification of a resistance dedifferentiation pathway and should be considered in combination with AR signaling inhibitors for ARPC with characteristics of lineage plasticity.

## INTRODUCTION

Targeted therapies are designed to attack cancer cells through specific molecular pathways to maximize impact and minimize general toxicity to the patient. Cancer cells can develop resistance to targeted therapies through a process of transdifferentiation where drug-sensitive tumor cells modify their lineage to acquire an alternate cellular identity that is not dependent on the targeted pathway for survival (*1-3*). Transition from an adenocarcinoma (ARPC) to neuroendocrine (NE) lineage is one differentiation path taken in various epithelial cancers including lung and prostate (*1, 4*). In metastatic castration-resistant prostate cancer (mCRPC), a decrease in luminal epithelial identity upon treatment with potent AR pathway inhibitors occurs in ∼20% of cases (*1, 5, 6*). Acquisition of a less differentiated, stem-like state leads to a spectrum of mCRPC phenotypes, termed lineage plasticity, which appears to exist along a continuum (*7*). A detailed characterization of stem-like cells across various models representative of heterogeneous CRPC are needed to gain insight into the process and to identify cellular points of therapeutic vulnerability. The stages of lineage plasticity most proximal to ARPC are thought to be CRPC with low AR signaling as well as CRPC representing an adenocarcinoma lineage that gains NE features while maintaining AR activity, so-called amphicrine (AMPC) subtype (*7*). The transition to highly aggressive small cell neuroendocrine prostate cancer (scNEPC) or a mesenchymal/basal AR^-^NE^-^ phenotype is thought to often proceed via clonal selection following the loss of *RB1* and/or *TP53* function (*2*).

The contribution of epigenetic mechanisms to cancer is a growing area of interest. Mutations to epigenetic regulators encompass functional oncogenes and tumor suppressors. BAF (also known as SWI/SNF) complexes remodel chromatin structure, playing a crucial process in gene regulation (*8*). Mistargeting of BAF complexes by disease-relevant transcription factors contributes to cancer initiation and progression. *ARID1A* and *ARID1B* are mutually-exclusive members of the cBAF complex, and *ARID1A* is mutated in ∼20% of cancers. *ARID1A* mutations in breast cancer determine resistance to estrogen receptor (ER) targeted therapy by promoting cellular plasticity resulting from loss of chromatin interactions for luminal lineage-determining transcription factors such as ER, FOXA1, and GATA3 (*9, 10*). In addition, *ARID1A* mutations occur with increased frequency relative to adenocarcinomas in neuroendocrine breast cancers (*11*) and are considered drivers for some neuroendocrine tumors, including gastrointestinal NETs and pulmonary carcinoids (*12, 13*). These data suggest that epigenetic deregulation of transformation is one mechanism associated with differentiated neuroendocrine tumors while small cell neuroendocrine tumors are driven mechanistically by *RB1* and *TP53* pathway mutations.

We established and characterized a patient-derived, CRPC organoid model harboring mutations in *ARID1A* and *ARID1B*, which demonstrates an amphicrine phenotype, consistent with *ARID1A* associated lineage plasticity Amphicrine CRPC has been increasingly recognized in patient samples and has been observed in the VCaP cell line. We describe a tractable model which has captured the phenotypic heterogeneity documented in the originating patient samples and provides a clinically-relevant model to study the dynamic interrelation of the ARPC and NE lineage states. Using lineage tracing, we experimentally determined the proliferative and differentiated status of subpopulations across ARPC, NEPC, and AMPC organoid subtypes. Following integration of single cell RNAseq clusters across multiple models, we observed the existence of a common stem/progenitor cell phenotype consistently present in all models, which is rare among AMPC and ARPC and frequent in scNEPC models. In ARPC there is a unique population of dividing, differentiated cells, while differentiated amphicrine cells are not self-sustaining but are continuously generated from progenitors. We also determined that chromatin accessibility in individual differentiated amphicrine cells is highly variable, suggesting appreciable stochasticity in the process of epigenetic lineage commitment secondary to ARID1A/B loss. Finally, we show that inhibition of stem/progenitor cells is sufficient to inhibit the growth of patient-derived AMPC.

## RESULTS

### Patient-derived organoids with mutations in BAF core complex components demonstrate lineage-plasticity and NE differentiation

We established a set of patient-derived organoid models designated NCI-PC35-1 and NCI-PC35-2 (*14*) (PC35-1 and PC35-2) from two spatially-separated needle biopsies of an mCRPC lymph node metastasis that was histologically ARPC with islands of NE marker-expressing cells (Figures 1A&B and S1A). There was no evidence of neuroendocrine markers in the primary tumor (Figure S1B). Whole-genome sequencing (WGS) phylogenetic analysis revealed that PC35-1 and PC35-2 arose from a common ancestor in the primary tumor featuring genomic mutations with high oncogenic potential: a deep deletion of *CDKN1B,* a frameshift mutation of *ARID1A* and a small deletion in *ARID1B* (Figures S2A and S2B). ARID1A/B are core components of the BAF complex, and reductions of ARID1A and ARID1B have been shown to drive carcinogenesis and neural developmental disorders (*15, 16*). The *RB1*, *TP53* and *PTEN* loci were intact and expressed (Figure S2C). Phylogenetic analysis showed little geographic co-mingling of PC35-1 and PC35-2, which demonstrated 77% and 97% exclusive subclonal genomic variants, respectively. Thus, the two PC35 models shared driver mutations and similar phenotypes but represented divergent clonal populations from separated core biopsies within the heterogenous tumor. We identified a tandem duplication of the AR enhancer in the metastatic samples but not the primary tumor (Figure S2D), and no additional known driver mutations were found in the metastatic clones, suggesting epigenetic regulation of lineage plasticity, consistent with mutations in the BAF complex. An analysis of prostate cancer cohorts in cBioportal and WCDT data sets revealed that *ARID1A* and *ARID1B* mutations occur at a frequency of ∼1 and ∼2% and demonstrate enrichment to ∼6 and 15%, respectively, in the CRPC setting, consistent with promotion of androgen resistance (Figure S2E).

**Figure 1.**
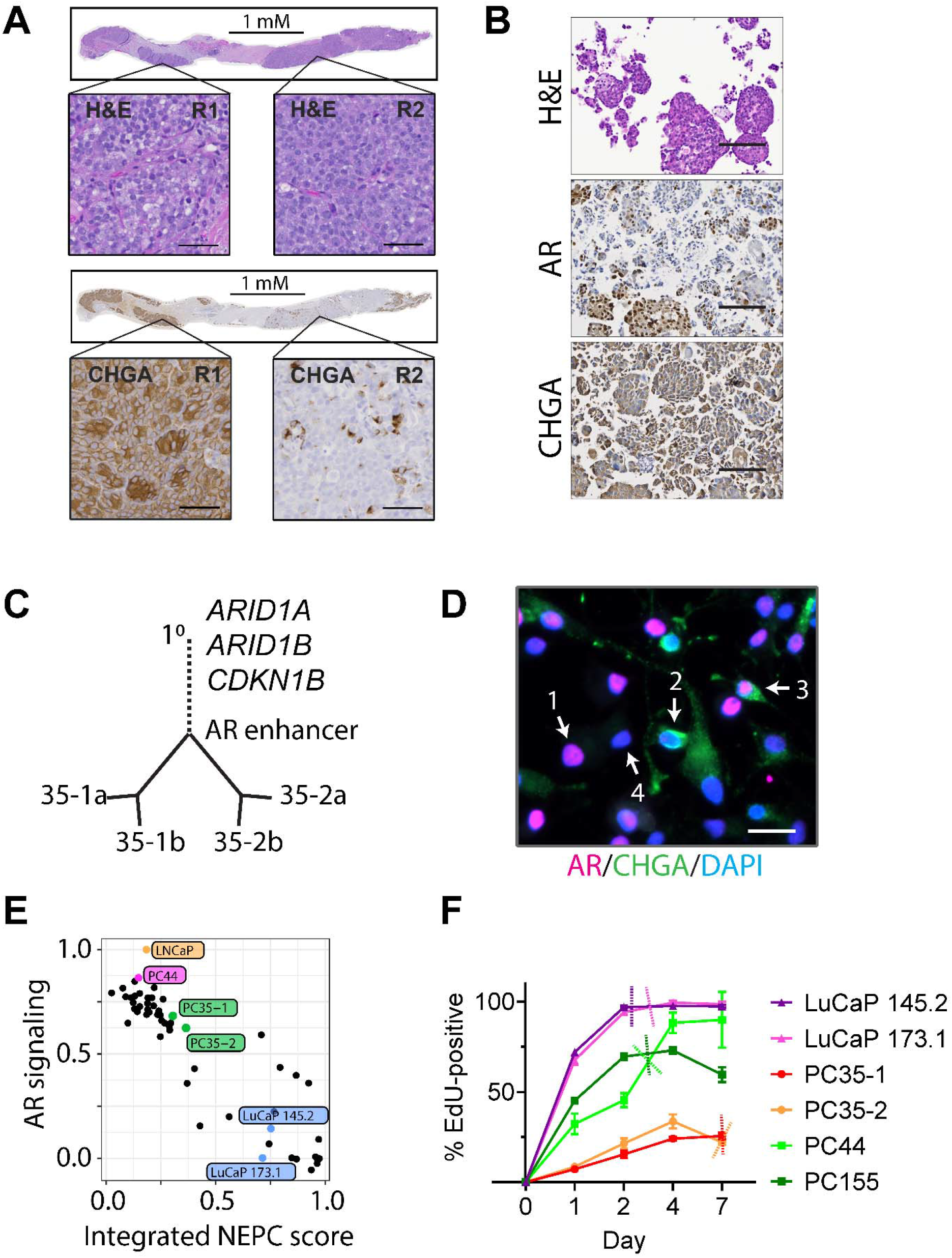
Patient-derived organoid models of mCRPC capture and maintain genetic and phenotypic heterogeneity. (A) Serial sections of tumor biopsy tissue stained with hematoxylin and eosin (H&E) or the indicated antibodies. Magnified views of Region-1 (R1) and Region-2 (R2) are shown. Scale bars, 50 μM. (B) Patient biopsy derived organoid sections stained with H&E or the indicated antibodies. Scale bars, 200 μM. (C) Phylogenetic tree of primary tumor and metastasis derived organoids. Primary prostate tumor, 1°. PC35-1 subclones - 35-1a, 35-1b. PC35-2 subclones - 35-2a, 35-2b. Significant genetic events indicated at positions in the tree where they originated. (D) PC35-1 organoids were dissociated to single cells and stained by immunofluorescence (IF) with antibodies against the indicated proteins and DAPI. Each of four phenotypes is indicated by a number and arrow. 1 = AR^POS^/CHGA^Lo/NEG^; 2 = AR^NEG^/CHGA^Hi^; 3 = AR^POS^/CHGA^Hi^; 4 = AR^NEG^/CHGA^Lo/NEG^. (E) The Weill Cornell Medicine cohort of mCRPC (black filled circles) and the indicated models used in this study are plotted by AR signaling score and NEPC score. (F) Continuous EdU-incorporation assay for the indicated organoid lines. The graph shows the percentage of EdU-positive cells over time. Dashed lines mark the approximate day of population doubling for each. Bar and line graphs are plotted as the mean of three independent experiments. Error bars represent ± standard error of the mean (SEM).

The organoids reflected the pathology of the metastatic tumor with a range of AR and/or NE-marked populations (Figure 1B&D). All possible combinations of AR and NE marker (CHGA) expression status were observed in individual cells. Further, mapping AR activity and NE signature scores of bulk RNA-seq data relative to other ARPC and NEPC models and clinical samples placed PC35-1/2 at the NEPC-adjacent edge of the ARPC cluster, consistent with the pathology and an early evolutionary step in ARPC lineage switching (Figure 1E). The distribution of phenotypic markers was stable over time (Figure S3A). EdU labeling and growth kinetic experiments showed that a minor population of PC35-1/2 cells underwent multiple divisions while the majority of cells were not proliferative (Figures 1F and S3B&C). This stands in contrast to scNEPC and ARPC organoid models, where a majority of the cells are proliferative (Figure S3C).

### Single cell RNAseq reveals subclonal heterogeneity and lineage gradients

To better understand the heterogeneity and subpopulation dynamics of the organoid models, we performed single-cell RNA-sequencing (scRNA-seq). By this method we identified two major clusters (I, II) in PC35-1 and three (I, II, and III) in PC35-2, with a heterogeneous range of lineage phenotypes (Figure 2A). Clusters were designated by phenotype based on AR, NE, and proliferation (PRLF) signature scores (Figures 2B-D and S4) as well as by comparison to a variety of phenotypically and biologically defined signatures for prostate cancer models, presented subsequently in Figures 4 and S9. Additionally, we performed RNA-velocity analysis (*17*) on the data to infer temporal states of differentiation (Figure S5). Stem/progenitor cells (St/Pr), which separated into two subclusters, colored red and blue, were designated based upon a high proliferation score, enrichment of stem marker gene expression, long RNA velocity vectors showing a multidirectional pattern (Figure S5), and a prominent G2/M transcriptional profile (Figure S4D). Prominent in both PC35-1 and 2 were an amphicrine population (colored orange) demonstrating features of both AR- and NE- gene expression, a low proliferation score, and a uniform direction of short RNA velocity vectors. Amphicrine populations demonstrated gradients of AR and NE-dependent gene expression (Figures S4B&C). In addition, PC35-2 had two additional ARPC clusters, one enriched for NR3C1-dependent transcription, and PC35-1 had an additional mesenchymal cluster. In contrast to PC35-1/2, the ARPC model PC44 and NEPC model LuCaP 145.2 were homogenous in their respective lineages and PRLF scores, indicating a more equal proliferative potential for the majority of cells in the population (Figures 2B-D). In summary, heterogeneity of PC35-1/2 captured from the patient tumor includes both inter- and intra-cluster lineage-plasticity.

**Figure 2.**
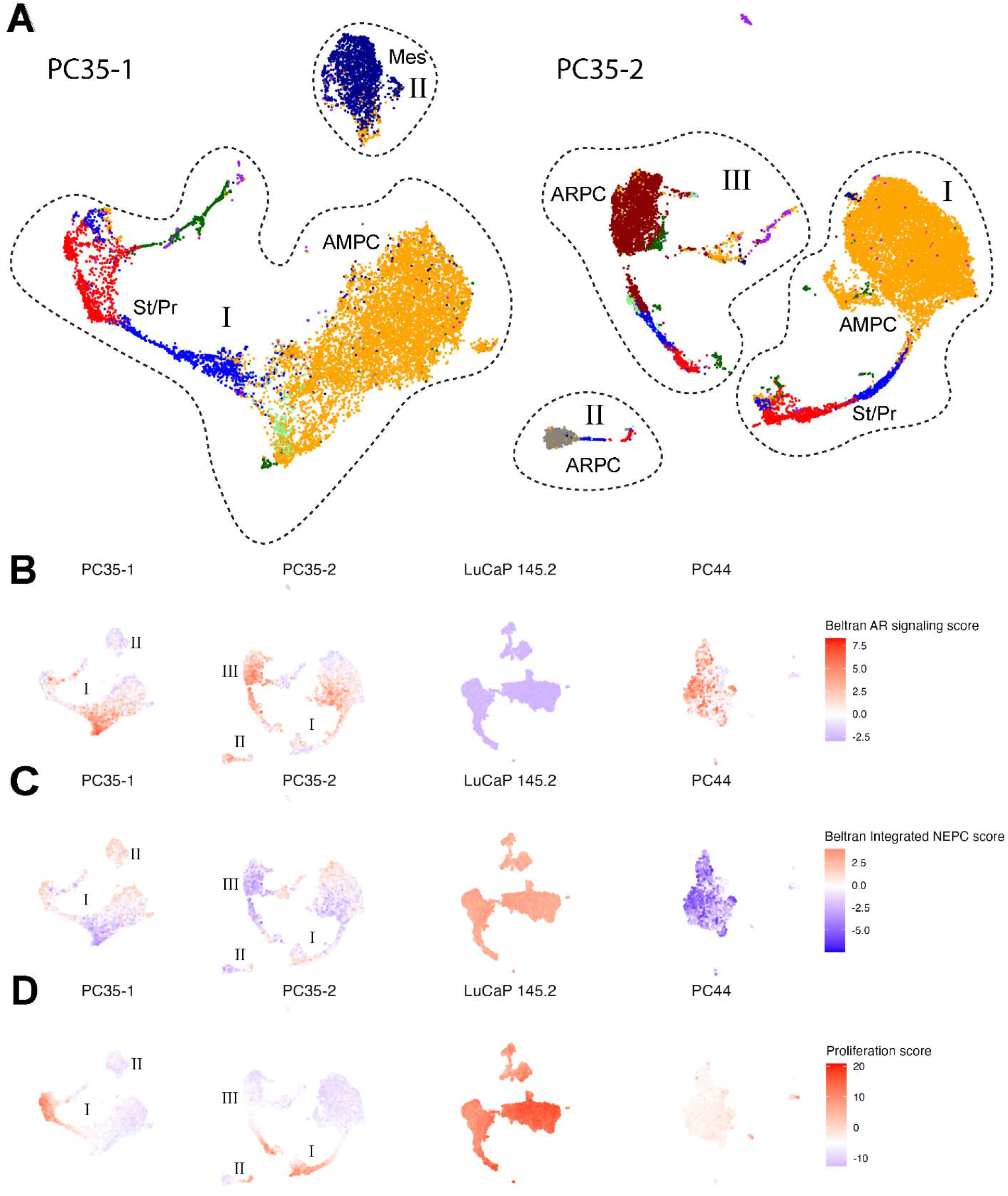
Single-cell transcriptomics identifies lineage-distinct heterogeneity in the PC35 organoid models. (A) scRNA-seq transcriptomic profiles of PC35-1 and PC35-2organoids plotted as UMAPs. Major clusters are circled and labeled with roman numerals (I, II, II). Clusters are colorized by phenotype designations, which are further characterized in Figures 4 and S9. Broad designations include stem/progenitor (St/Pr) = red and blue, adenocarcinoma (ARPC) = brown (NR3C1-enriched) and grey, amphicrine (AMPC) = orange, or mesenchymal (Mes) = dark blue. (B) AR and (C) neuroendocrine (NE) signature scores for each cell determined by principal component analysis (PCA) using published gene sets from Beltran et al. Loadings from the first principal component for each cell are projected onto the UMAPs from (A) and UMAPs plotted for LuCaP 145.2 and PC44 scRNA-seq transcriptomic data. (D) Proliferation score determined as in (B) and (C). The proliferation gene set was derived from Balanis et al.

**Figure 3.**
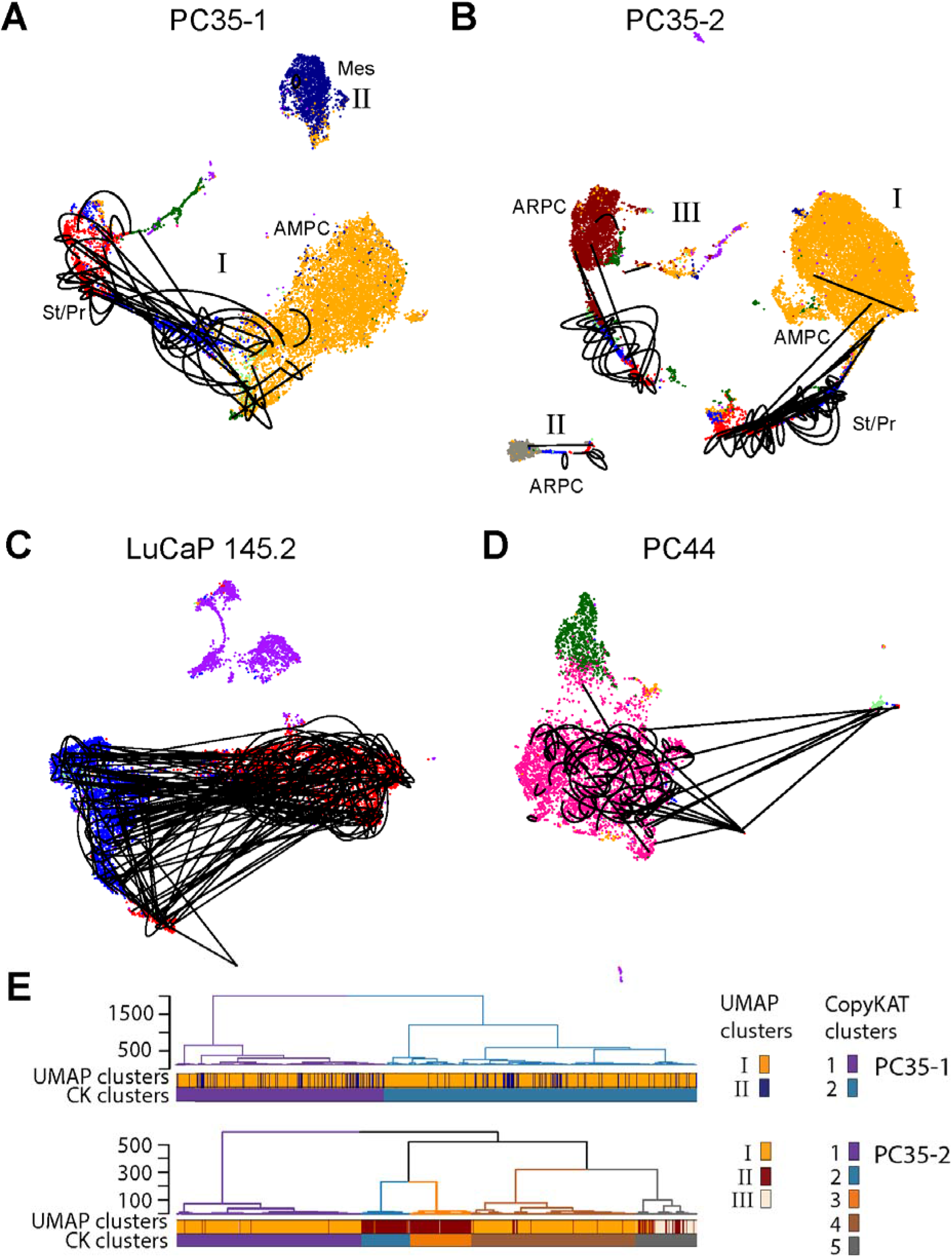
Single-cell combinatorial barcoding identifies lineage-distinct and stem- like/progenitor subpopulations. (A-D) CellTag lineage-tracing analysis. Clusters are colorized by phenotypes further defined in Figures 4 and S9. (A) UMAP major clusters I and II of PC35-1 and (B) I, II, and III of PC35-2 are shown. Each cell in a clonal population (≥ 2 cells expressing the same combination of barcode IDs), is connected by a black line. Self-renewing clones that exist in the same subcluster are connected by curved lines, differentiating clones that span at least two subclusters are connected by straight lines. (C) Clonal connections as in (A) and (B) mapped onto (C) LuCaP 145.2 and (D) PC44 UMAPs. (E) Comparison of subpopulations defined by their scRNA-seq transcriptional profile (UMAP clusters), to subclones defined by genomic CNV (CopyKAT clusters) determined using the same scRNA-seq data. Results for both PC35-1 (top) and PC35-2 (bottom) are shown. Dendrograms are colored according to CopyKAT cluster number and show the hierarchical relationships among the CopyKAT clusters. Heatmaps directly below the dendrograms show the distribution of cells from UMAP major clusters I, II, and III throughout the CopyKAT clusters. Each cell is represented by a vertical line colored according to the UMAP cluster (top row) or the CopyKAT cluster (bottom row) to which it belongs and sorted by CopyKAT cluster. PC35-2 contains two additional minor UMAP clusters lacking differentially-expressed genes that were not annotated here.

**Figure 4.**
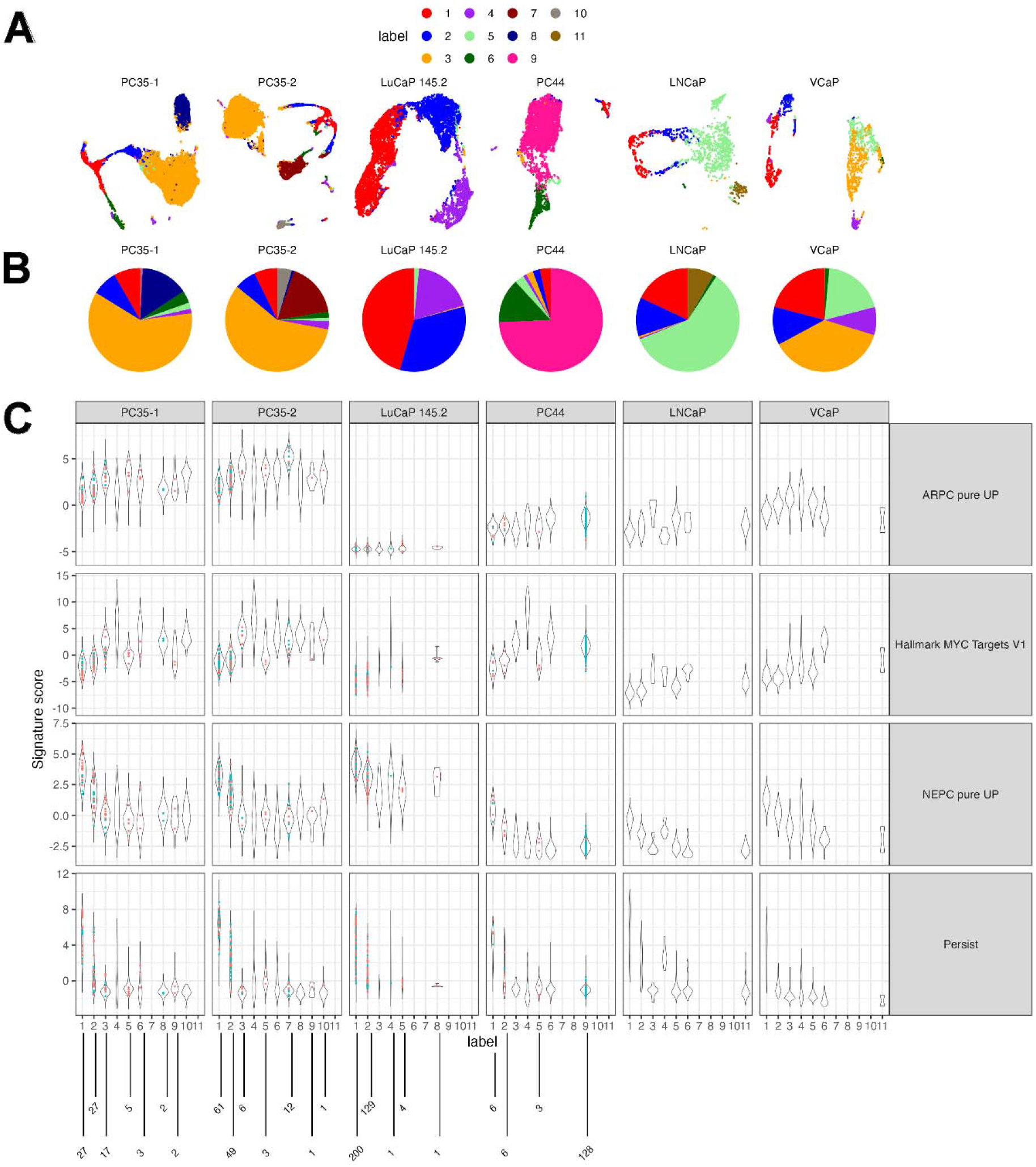
Integration of organoid cohort with LNCaP and VCaP cell lines shows common phenotypes across samples. (A) UMAP embeddings of integrated scRNAseq for NCI-PC35-1/2, LuCaP145.2, and NCI-PC44 organoid cultures and LNCaP and VCaP cell lines using mutual nearest neighbors correction. Each UMAP was generated using the batch corrected principal components for each sample independently. Cells are colored based on clustering membership using integrated data for all samples. 1) Stem/progenitor (St/Pr)-1, 2) St/Pr-2, 3) AMPC, 4) NEPC, 5) ARPC/luminal stem (*41*), 6) ARPC/NR3C1^HI^, 7) AMPC, 8) mesenchymal, 9) ARPC/NR3C1^HI^, 10) AMPC, 11) ARPC/luminal stem, (B) Pie charts show the relative proportion for each cluster in each sample. (C) Violin plots show signature scores relative to cluster and model. Colored dots are signature scores for individual cells that were determined to be CellTagged clones. Red dots are self-renewing clones, blue dots are differentiating clones. The number of clones is enumerated below each cluster.

To validate the scRNA-seq analysis and to develop a quantitative assay for specific subpopulations, we performed quantitative single-molecule RNA-FISH using selected markers on PC35-1/2 organoid-derived cells (Figure S6). In agreement with the scRNA-seq results, we found ARPC lineage marker-positive cells, NEPC marker-positive cells and double positive cells (Figure S6A). EZH2 and TK1 marked the same population of EdU-positive, dividing cells (Figures S6A-C). Double staining for mRNA and protein of selected lineage markers confirmed a strong positive correlation between mRNA and protein (Figure S6D).

### Simultaneous tracking of lineage and clonal identity with single-cell resolution identifies self-renewing stem-like subpopulations with differentiation potential

To address the dynamic plasticity and the hierarchical structure of the subpopulations, we used the “CellTagging” method of combinatorial indexing of expressed barcodes, determined by scRNA-seq, to simultaneously track cellular origin and phenotypic identity within growing organoids (*18*). We analyzed single time-point experiments of different durations after tagging to identify clonal expansion of sibling cells, allowing determination of the clonal relationships for lineage-marked populations. A separation of four weeks between tagging and harvest captured uniquely tagged clones in all major clusters. For both PC35-1 and PC35-2, all tagged clonal sibling cells that were associated with any given major cluster were exclusive to that cluster, demonstrating the clonality of each cluster. The location of tagged sibling cells is graphically depicted on the UMAPs for PC35-1/2 with black lines connecting clones (Figures 3A&B). Within each major cluster a disproportionate number of the tagged clones were located entirely within the stem/progenitor subclusters (quantified in Figure S7), indicating self-renewal. Tagged stem/progenitor clones also spanned across the differentiated subclusters within the same major cluster, demonstrating differentiation. The more differentiated subclusters showed very limited internal cellular replication, suggesting that differentiation from a cancer stem/progenitor cell was the major source of amphicrine cells.

In contrast, the clones captured in LuCaP 145.2 NEPC organoids were indicative of a widely proliferative population. We detected numerous clones both within and nondirectionally across the two major phenotypically identified stem cell populations, while the small neuroendocrine population (colored purple) contained no tagged clones (Figure 3C). The ARPC model PC44 exhibited two phenotypes of dividing cells, a high number of clones associated with two small clusters and giving rise to differentiated daughters as well as self-renewing cells within the main luminal cluster (Figure 3D). In contrast, AR^+^ cells derive mainly from stem cells in the AMPC model.

One possible explanation for the discrete clonality of the major clusters in the PC35 models is that cluster-specific genetic events led to distinct phenotypes, although we were unable to identify subclonal driver mutations by WGS analyses. We analyzed our scRNA-seq data using CopyKAT to identify clonal subpopulations based on genomic copy number variation (CNV) and associated this genomic substructure with the phenotypically-defined major clusters (*19*). We found two patterns defined by contributing CNV-identified clones to the different phenotypes of the clusters: those which were independent of clonal patterns (AMPC, mesenchymal, and PC35-1 ARPC clone II phenotypes) and those attributed to unique or closely related clonal genotypes (the PC35-2 ARPC III clone) (Figures 3E and S8A&B). These data suggest that there are fixed differentiation patterns for pre-existing clonal populations in addition to common pathways of lineage plasticity, demonstrating multiple contributing factors to phenotypic heterogeneity.

### Integration of single cell RNAseq across CRPC models suggests distinct growth patterns associated with lineage subtypes

To compare tumor subpopulations among ARPC, AMPC, and NEPC, we integrated scRNAseq data across the organoid models and in addition included published scRNAseq data from two commonly-used metastatic cell lines, LNCaP and VCaP (Figures 4A&B). We also mapped individual, tagged daughter cells onto the integrated clusters (Figure 4C). The distribution of clusters among the models and their relative expression of AR, NE, MYC, and Stem/Progenitor (St/Pr) signatures is shown (Figure 4C). Additional lineage and relevant biological signature enrichments are shown in Figure S9.

All models contained stem/progenitor cell populations, which separated into two subclusters. St/Pr 1 (colored red) appeared to be the originating stem population, while St/Pr 2 (colored blue) demonstrated transcriptomic properties of a transit amplifying subpopulation, as well as defining the cluster proximal in similarity to differentiated cells. While the St/Pr populations were relatively minor in the ARPC and AMPC models, they composed the majority of the scNEPC cells (Figure 4B).

The St/Pr signature is overlapping with a signature termed ‘Persist’, which was identified in LNCaP cells from an enzalutamide resistant subpopulation, and computational approaches inferred ‘Persist’ as a founding trajectory population (*20*). Similarly, in primary prostate cancers, the ‘Sig51’ signature, which was shown to correlate with increased aggressiveness, overlaps the integrated St/Pr signature (Figure S9) (*21*). By contrast, the gene set defining the recently-described, mesenchymal stem-like PC subpopulation (termed MSPC) in bulk cultures of mCRPC cell lines, organoids, and xenografts (*22*) is not substantially enriched in the St/Pr signature defined here. In addition, using experimental lineage tracing, we show here for the first time in CRPC models a direct one to one correspondence in individual cells between either self-renewal or the generation of differentiated daughter cells and the expression of a gene signature (St/Pr), which has been associated with enzalutamide resistance and clinical aggressiveness.

The ARPC cell line models share a major population (cluster 5, enriched for genes found in mouse basal and castration-resistant luminal cells), which is a minor population in the organoid models (Figures 4A&B). However, in the organoid models, cluster 5 cells are juxtaposed or intermixed with St/Pr-2 cells, and in some cases, demonstrate evidence of division, i.e. lineage tracing (Figure S7). One possibility is that cluster 5 cells represent an intermediary cell that may have been expanded in the continuous growth of cell lines selected on a two dimensional surface. Amphicrine cell populations, which have yet to be thoroughly characterized due to their rarity in tractable models, are found in the NCI-PC351/2 and in the VCaP cell line (Figures 4A&B).

Signatures characterizing amphicrine populations (clusters 3,7,8, and 10) include senescence, mesenchymal, ‘Prosgenesis’ (regenerative luminal cells enriched for basal and mesenchymal features) (*23*), MSPC (*22*), and clusters from LNCaP models that have been selected by long-term growth in enzalutamide (Figure S9) (*20*). This suggests that amphicrine transcriptomes are a component of various cell line and mouse models describing progression following AR signaling suppression of ARPC.

### Single cell ATACseq reveals transcription factors regulating amphicrine stem cell, luminal, and mesenchymal pathways

To characterize transcriptional regulation in an epigenetically driven amphicrine model, we performed single-cell ATAC-seq on PC35-1/2 organoids. Clustering by genome-wide chromatin accessibility yielded three clusters (***1****, **2**, **3***) in both PC35-1 and PC35-2 (Figure 5A). To look for transcription factors (TFs) that may be responsible for the differing phenotypes among the clusters, we performed an analysis of inferred TF activity and assessed the similarity with ATACseq-based signatures describing various subclasses of clinical mCRPC or dynamic states of lineage transition (*22, 24*) (Figure 5B-D).

**Figure 5.**
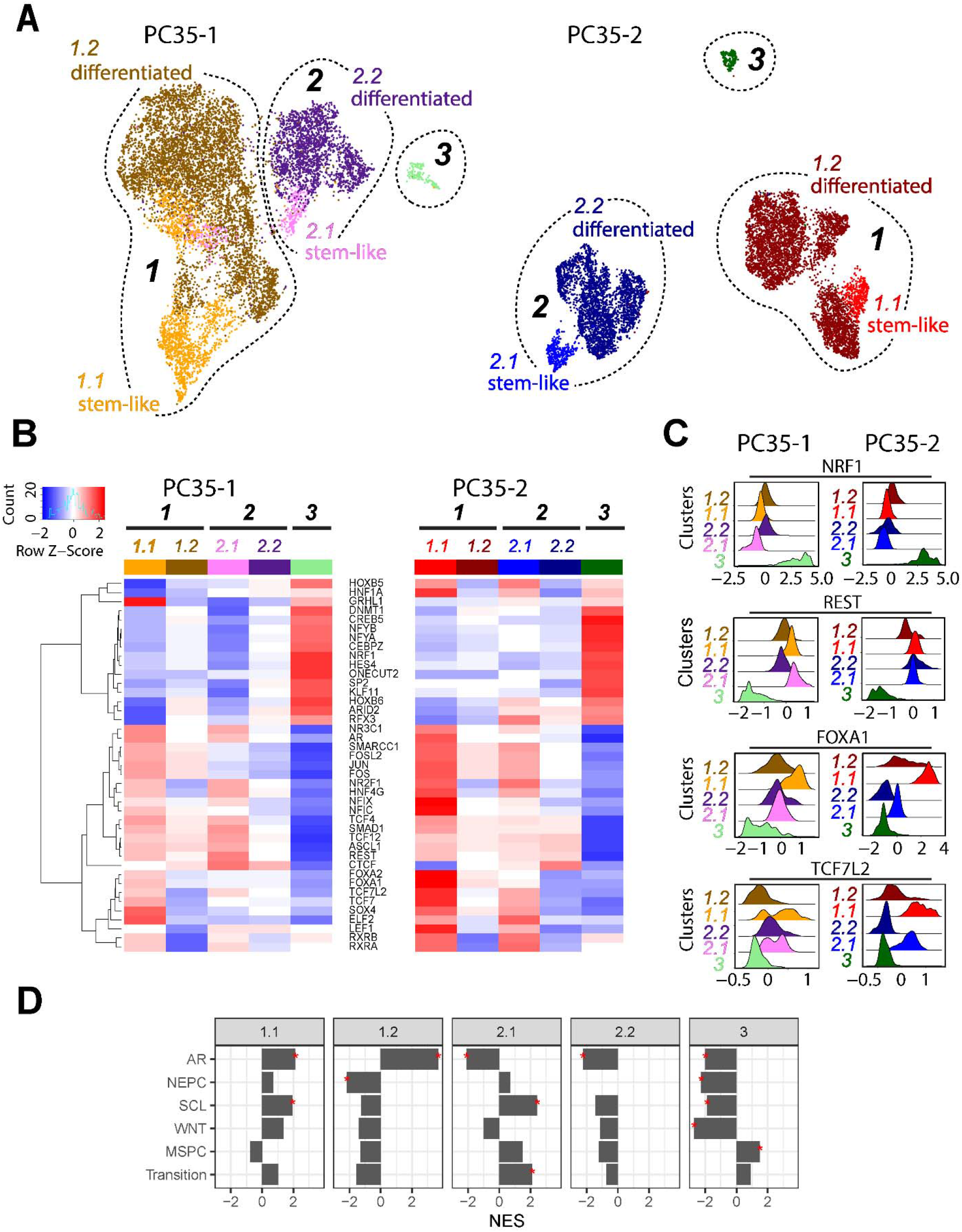
scATACseq clusters NEPC and ARPC populations in the PC35 organoids. (A) UMAPs of global chromatin accessibility for PC35-1 and PC35-2. Major clusters are annotated as *1*, *2*, *3*. Clusters *1* and *2* are partitioned into two additional subclusters, stem-like and differentiated. (B) Heatmaps show inferred transcription factor (TF) activities of the listed TFs for each of the UMAP clusters/subclusters in PC35-1 and PC35-2. The heatmaps are colored by deviations z-scores, calculated as the differential accessibility for a particular transcription factor compared to the average accessibility profile for the entire dataset. TF activity shown was selected using the score markers function (see Methods) on the deviations z scores to identify maximal enrichment across lineages and development. Transcription factors shown were determined to be expressed and selected from a list of the top fifty most deviant TFs. (C) Inferred transcription factor activity density plots for each cluster population. Deviations z scores are shown on the x-axis for the TF indicated at the top. Density estimates are represented along the y-axis and broken down by cluster/subcluster. (D) Normalized enrichment scores derived using GSEA for each cluster across molecular phenotyping signatures. Significant scores are denoted with “*” (adjusted P ≤ 0.05).

Clusters-***3*** in both models were distinguished as mesenchymal-like, lacking stem cell (SC), ARPC, NEPC, and WNT driven phenotypes (Figure 5D). These clusters were notable for a relative absence of REST activity and high activity scores for TFs such as NRF1, HES4 and ONECUT2 (Figures 5B&C).

Clusters ***1*** and ***2*** in PC35-1/2 demonstrated unique but highly overlapping combinations of transcription factors contributing to stem cell, luminal epithelial, and neural phenotypes. Additionally, Clusters ***1*** and ***2*** of both models could be partitioned into two pairs of subclusters *1.1*, *1.2* and *2.1*, *2.2* (Figure 5B). Inferred TF activities in subclusters *1.1* and *2.1,* were consistent with a stem-like phenotype, while 1.1 and 2.1 were enriched for ATAC signatures determining luminal epithelial and transitional mesenchymal lineages, respectively (Figures 5D). The “transition” signature includes JAK/STAT inflammatory processes and bridges stem cell and neural populations in a mouse model (*25*). Consistent with the expression of NE markers in a mouse model transitioning from adenocarcinoma to NEPC, ASCL1 and FOXA2 transcription factor activities were observed, but other classical scNEPC transcription factors such as MYCN, BRN2 or NEUROD1 were not apparent (*26-28*). Of interest, a WNT related signature was exclusively expressed in the luminal epithelial stem cells but lost in the transitional mesenchymal population. These data extend and contextualize previously-defined CRPC subclass signatures to plasticity-mediated subpopulations.

There were no remarkable TF activities gained in Clusters *1.2* and *2.2* compared to the stemlike *1.1* and *2.1.* The defining signatures in 1.2 were enriched AR-driven and depleted NE signatures, while 2.2 demonstrated a depleted AR signature. Of interest, differentiation was mostly associated with reduced TF activity relative to the stem-like clusters. One possible explanation for this is that the plasticity-associated heterogeneity across the differentiating population obscured TF patterns. This is supported by the observation that there is a loss of accessibility for the TFs shown in Figure 5B, while the average global number of accessible sites did not change. These data suggest a model of plasticity whereby activity of a transitional mesenchymal lineage program on luminal stem-like/progenitor clones resulted in distinct cellular phenotypes that share to differing degrees features of ARPC and NEPC lineages, as observed in the major amphicrine cellular population.

### Targeting both AR pathway dependent and independent compartments of the stem/progenitor subpopulations inhibits *in vitro* and *in vivo* tumor growth

The existence of multiple identifiable clones propagated almost exclusively by stem/progenitor cells in the AMPC model implies an approach to treatment. Because at least a portion of the St/Pr population expressed AR, we incubated PC35-1 and PC35-2 organoids with enzalutamide, quantified cell numbers, and found a partial response in both models, concordant with the notion of a subpopulation-specific dependence on AR signaling (Figure S10A). PC35-1 showed a greater than two-fold reduction after treatment, while PC35-2 showed a less than 30% decrease. We then performed RNA-FISH in combination with EdU to quantify subpopulation-specific changes due to enzalutamide treatment (Figures S10B-D). Congruent with the different overall response observed in bulk, we found that enzalutamide caused a >10-fold reduction to proliferating AR^POS^EdU^POS^ cells in PC35-1 while the same population in PC35-2 showed only a small decrease (Figure S10C). PC35-2 demonstrates an NR3C1-dependent signature, consistent with enzalutamide resistance (PC35-2 cluster 7, Figure S9). The SCG2-positive, neuroendocrine-like, populations in both PC35-1/2 were insensitive to enzalutamide (Figure S10D) (*29*). These data document clonal variability, selected within a patient, leading to partial ARPI resistance within a population of mCRPC tumor cells with no known differences in oncogenic drivers.

Although AR^POS^ cells made up a proportion of the EdU^POS^ progenitor population, **≥** 50% of stem/progenitor cells were AR-negative (Figure S10C, -Enza columns). To specifically address the stem/progenitor population, we identified multiple druggable targets as highly enriched in the St/Pr clusters, including EZH2, AURKA, and the Notch pathway (Figures S4B&C) and targeted them with CPI-1205, Alisertib, or Compound E, (EZH2i, AURKAi and Notchi respectively). For comparison, we included the chemotherapeutic agent carboplatin, which is used as a late line of therapy in mCRPC. We treated the PC35 organoids with AURKAi, EZH2i, Notchi, carboplatin, or DMSO for six weeks. In both organoid models, the AURKAi caused a nearly 10-fold decrease in cell number compared to DMSO while the other drug conditions resulted in only minor reductions (Figure 6A). We tracked the effect of AURKAi, EZH2i, and carboplatin relative to DMSO with single-cell resolution using RNA-FISH/EdU combined assays. Subpopulations were identified by marker gene expression: AR to mark ARPC lineage; SCG2 to mark NEPC lineage; TK1, EZH2 and AURKA to mark stem-like/progenitors. We found that the AURKAi specifically depleted the dividing, EdU-incorporating stem-like/progenitor subpopulation in PC35-1/2 (Figure 6A&B), while the SCG2 population increased as a percentage of the total (Figure 7B&C). By comparison to AURKA inhibition, carboplatin treatment of PC35-1/2 had no effect (Figure 6A-C).

**Figure 6.**
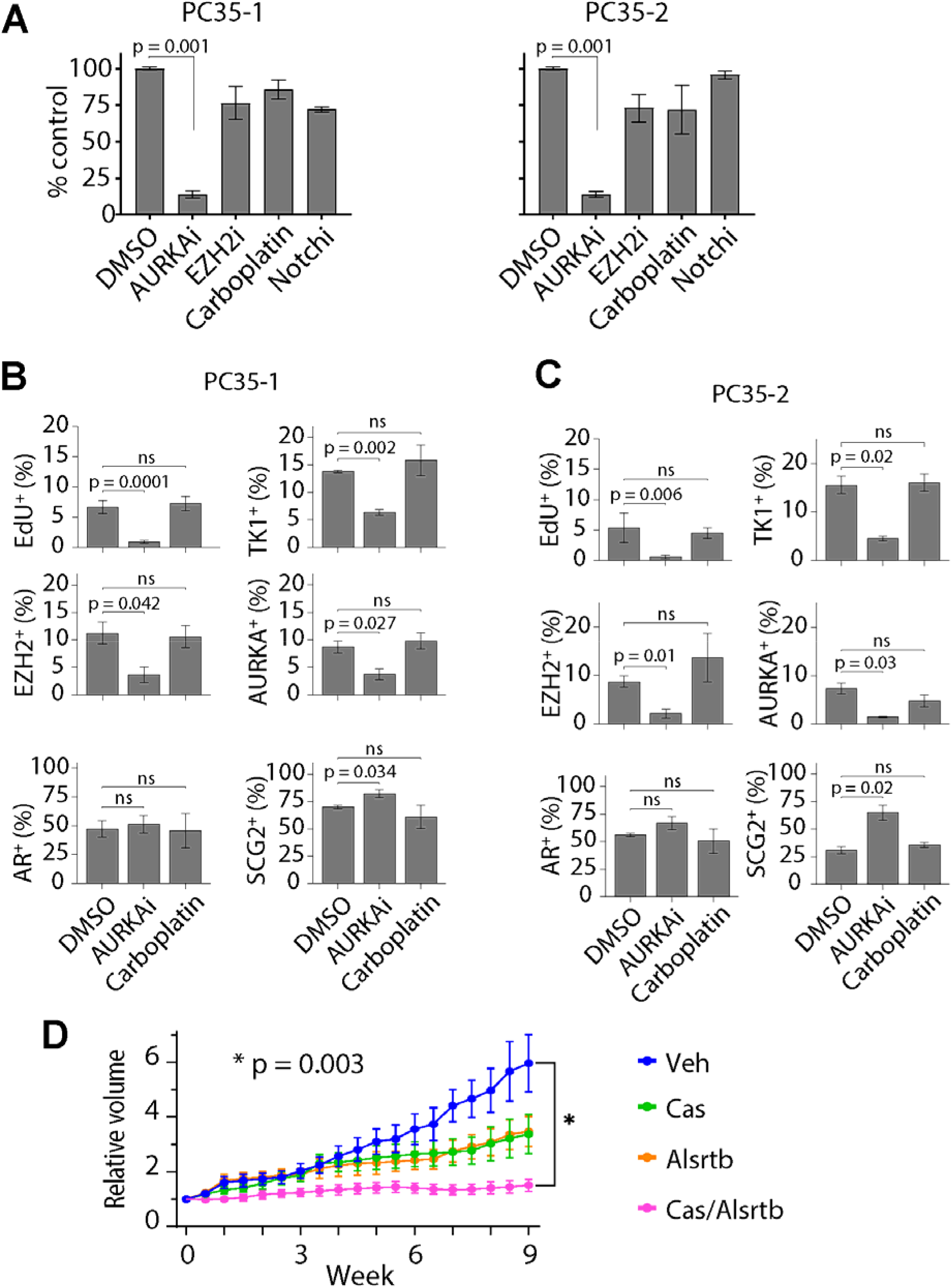
The stem-like/progenitor subpopulation is vulnerable to AURKA inhibition. (A) Determination of drug sensitivity. Organoids were treated twice weekly for six weeks with 500 nM AURKAi, 500 nM EZH2i, 500 nM carboplatin, 1 μM Notchi, or 0.02% DMSO-treated controls. Quantification was done by dissociating the organoids and manually counting the cells. The quantified values for each condition were plotted relative to the DMSO controls. (B) PC35-1 and (C) PC35-2 organoids treated for six weeks as in (A) with AURKAi, carboplatin, or DMSO, then pulsed with 10 μM EdU, 24 hours prior to collection. The indicated marker expression for each cell was determined by RNA-FISH. EdU incorporation status (positive or negative) was determined for each cell. Data was plotted as the percentage of cells that were positive for a given marker or EdU for the three treatment conditions. (D) Relative change in tumor volume for PC35-1 organoid-derived xenografts (ODXs) during nine weeks of the indicated treatments. Tumor volume was calculated as an average of the replicates. The change in volume was calculated relative to the “0” time-point (∼150mm^3^ tumor volume). Vehicle n = 5 mice; castration n = 5 mice; alisertib n = 4 mice; castration + alisertib n = 5 mice. Bar graphs are plotted as the mean of three independent experiments. Error bars, ± SEM. P-values were calculated using the student’s t-test, two-tailed, unpaired.

To evaluate how effective inhibition of AURKA and/or AR is at blocking tumor growth *in vivo*, we treated PC35-1 organoid-derived xenograft tumors for nine weeks with either alisertib, castration, alisertib combined with castration, or vehicle. Castration or alisertib (30 mg/kg once/day) alone caused a 50% decrease of tumor growth. However, the combination treatment rapidly and dramatically blocked tumor growth (Figure 6D). In addition, an alternative dose schedule of alisertib (20 mg/kg twice/day) alone, designed to maintain a continuous efficacious exposure level, caused tumor regression (Figure S11A). We validated the loss of dividing cells and retention of differentiated amphicrine cells in the residual tumor bed (Figure S12). In nine week CT35 tumors, castration caused a strong increase in cytoplasmic and decrease in nuclear AR, as well as increased expression of the NE marker, synaptophysin (Figure S12A). Consistent with the effects on tumor growth, the strong BrdU incorporation observed in the control was decreased in all the treated conditions, reaching the lowest level in the combination treatment (Figures S12A&B). These results demonstrate that the subpopulation-specific vulnerabilities that we identified in patient-derived organoids can be exploited to yield impactful results on tumor growth *in vivo*.

We analyzed combination drug responses in tumors from the other available AMPC model, the VCaP cell line (Figure S11B). VCaP tumors demonstrated strong castration sensitivity, consistent with virtually all tumor cells expressing AR, unlike PC35 that includes a notable AR^LOW/NEG^ population (Figure S11B). In contrast to PC35, alisertib alone was minimally active in VCaP xenographs. Although the rapid development of resistance to alisertib due to the mutational background of the model is one explanation for this difference, another possibility is suggested by the observation in the PC44 ARPC model, which contained dividing differentiated luminal progenitors, distinct from the common St/Pr population in PC35 and other models.

## DISCUSSION

There is a growing appreciation for the role of lineage plasticity in drug resistance and progression. Amphicrine (AR^+^NE^+^) CRPC tumors are thought to be an early indicator of lineage plasticity and are of particular interest in providing models to investigate mechanisms associated with lineage losses and gains. The amphicrine model investigated here is driven by mutations in the epigenetic regulators, *ARID1A* and *ARID1B,* and the cell cycle regulator, *CDKN1B,* and exhibited at the single-cell level all permutations of luminal epithelial and NE lineage marker expression. Although clinical amphicrine CRPC has been characterized for pathological and molecular features, tractable cell models have been limited to the VCaP cell line. Integrating scRNAseq data across multiple models and applying various CRPC lineage signatures revealed that amphicrine populations were enriched for mesenchymal stem cell, regenerative, and enzalutamide resistance characteristics, which are often assumed to be linked with progression (*20, 23, 25*).

We show the existence of clonally-distinct cancer stem/progenitor subpopulations as the source of growth and phenotypic heterogeneity. Surprisingly, the least proliferative population was the most neuroendocrine-like. This finding is contrary to the increased growth rate of AR^-^ NEPC driven by *RB1* and *TP53* loss, but consistent with less aggressive NE tumors including gastroenteropancreatic NE neoplasms, breast cancer with neuroendocrine differentiation, and pulmonary NE carcinoids that are frequently driven by mutations in *ARID1A* (*11-13*). This model is evidence for the concept that stem cell plasticity per se does not correlate with aggressiveness but that additional drivers are needed to overcome proliferative arrest associated with terminal differentiation. To the best of our knowledge, this is the first near-patient CRPC organoid that dynamically produces a range of phenotypically heterogeneous daughters. Additional models of amphicrine CRPC are needed to obtain a more complete understanding of the range of biological characteristics.

Diversified and labile transcriptional programs within a heterogenous tumor cell population can rapidly confer clonal fitness in the face of therapeutic pressure (*20, 30-32*). In addition to lineage-switching, we found ARPI resistance in independent clones mediated by other potential mechanisms, such as high expression of *NR3C1.* This observation directly demonstrates within a patient CRPC tumor that multiple different paths to AR resistance exists within the same tumor, underscoring the challenges in the development of curative treatments.

Despite the range of lineage heterogeneity, scATACseq identified two major populations of amphicrine stem cells with overlapping inferred transcription factor activity. Two populations were distinguished by ATAC signatures representing respectively either ARPC and WNT pathway or transitional JAK/STAT and mesenchymal pathways. The latter signature is kinetically associated with cells in a GEMM model that bridge adenocarcinoma and NE population transitions (*25*), suggesting a contribution in PC35 amphicrine cells to the expression of NE markers such as CHGB. Of interest, the differentiated populations had equally accessible global chromatin relative to the stem cell state but reduced inferred activity for a variety of the transcription factors. One interpretation of this data is that plasticity is associated with a widely-accessible chromatin state followed by heterogeneous and perhaps stochastic chromatin closing during commitment.

The existence of a stem-like/progenitor subpopulation as the seedbed of growth in a tumor predicts excellent potential as a point at which to direct therapeutic intervention. We found Aurora Kinase A, a regulator of mitotic progression, stem cell self-renewal, and asymmetric division (*19, 33, 34*), to be expressed and restricted to the stem/progenitors. Inhibition of AURKA in the organoid models caused a strong and specific depletion of the stem/progenitor pool that blocked growth of the entire heterogeneous population and importantly, blocked tumor growth *in vivo*. On the other hand, alisertib alone did not block VCaP growth, although castration was efficacious as was castration in combination with alisertib, as expected. The lineage hierarchy/origin of the amphicrine subpopulation in VCaP ARPC is unknown. In distinction to CT35, the VCaP cell line has been selected in culture for growth on plastic and expresses AR in almost all cells. Using lineage tracing in combination with scRNAseq, we observed in the PC44 ARPC model, the existence of two pools of dividing tumor cells, a minor St/Pr population and self-renewing luminal epithelial population. Thus, we anticipate that alisertib alone would likely not be useful in most ARPC tumors but may be beneficial in combination with AR pathway inhibitors to suppress the dividing St/Pr population that is insensitive to AR inhibition. Clinical trials targeting treatment of scNEPC with alisertib have been mostly ineffective. Our analysis of the population structure and its relationship to cell division potentially provides insight. >70% of the scNEPC cells had a St/Pr phenotype. Besides the underlying physiological differences attributable to distinct genomic mutations in scNEPC such as *RB1* loss, it is expected that targeting a major as compared to a minor population of St/Pr tumor cells would increase the probability of resistance. In total the results presented here establish the utility of alisertib for a cancer stem cell-driven model of AMPC and suggest that alisertib be tested as a combination agent for ARPC to assist with the development of lineage plasticity-mediated resistance to AR pathway inhibitors.

## ACKNOWLEDGMENTS

The authors wish to express their gratitude to the patients and the families of the patients who contributed to this study. We would like to thank the LGCP Microscopy Core at the NCI/CCR and we would like to thank the CCR Single Cell Analysis Facility. Sequencing was performed with the CCR Genomics Core. This work utilized the computational resources of the NIH HPC Biowulf cluster (http://hpc.nih.gov). We would like to thank A. Zoubeidi for providing the EZH2 phospho-T350 antibody. We thank D. Takeda, G. Merlino, J. Shern, and M. Shen for reviewing the manuscript.

## FUNDING

This research was supported by the Intramural Research Program of the NIH, National Cancer

Institute, Center for Cancer Research.

Prostate Cancer Foundation (Young Investigator Awards to M.L.B. and A.G.S.)

Department of Defense Prostate Cancer Research Program (W81XWH-16-1-0433 to A.G.S)

Support from CCR Single Cell Analysis Facility was funded by FNLCR Contract

HHSN261200800001E.

## AUTHOR CONTRIBUTIONS

Conceptualization: M.L.B., K.K.

Methodology: M.L.B., S.A.M., S.A., F.H.K., W.L.D., K.K.

Investigation: M.L.B., R.L., A.G.S., D.B., C.M.T., C.T., J.K., J.Y., A.N.A.

Formal analysis: M.L.B., B.J.C., A.T.K., K.K., R.L., A.G.S., J.Y., T.L.L.

Visualization: M.L.B., B.J.C., A.T.K.

Project administration: M.L.B., K.K.

Supervision: K.K.

Writing: M.L.B., K.K.

## MATERIALS AND METHODS

### Histology

Formaldehyde-fixed tissue and organoid sections were embedded in paraffin blocks. Sections were cut, mounted on slides, and put through steps of graded alcohol deparaffinization. Steam antigen retrieval was performed for fifteen minutes (DAKO 1699) followed by washes in PBS/0.1% Tween-20 (PBST) 3x five minutes. Sections were blocked in Background Buster (Innovex NB306) for 40 minutes and then incubated overnight in primary antibody at 4°C. The next day the slides were washed three times with PBST and then incubated with a biotinylated secondary antibody for 30 minutes at room temperature. Antibody staining was developed with 3, 3’ diaminobenzidine (DAB) and counterstained with hematoxylin. Slides were imaged using a Zeiss Axioscan.Z1 microscope with a plan-apochromat 20x NA 0.8 objective.

ODX tumor sections were processed and imaged as described above. The sections were stained using an Intellipath FLX autostainer (Biocare Medical). Quantification of BrdU was done using the Indica Labs HALO v3.3 software running the CytoNuclear v2.0.9 algorithm. The optical density threshold for “weak” labeling was 0.377 “strong” was set at 1.331. Data was plotted using GraphPad Prism v8.

The IHC for the biopsy tissue sections was done in the NIH Clinical Center Pathology lab.

### Organoid culture

Organoids were established and cultured according to our previously described methods and culture conditions (*14*). Patients provided informed consent, and samples were procured from the NIH Clinical Center under NIH Institutional Review Board approval in accordance with U.S.

Common Rule. NCI-PC35-1, NCI-PC35-2, and LuCaP 145.2 organoids were grown in PrEN - p38i/-NAC media conditions. NCI-PC44 organoids were grown in PrEN -p38.

### DNA extraction

DNA was extracted from the primary prostate tumor. Sections at 5 μM thick from the paraffin block of radical prostatectomy tissue were cut onto slides but not mounted and then stained with H&E. Tumor tissue from five sections was macrodissected and combined into one tube. Adjacent normal prostate tissue from 19 unmounted sections was combined into another tube. DNA was extracted using the QIAmp DNA FFPE Tissue Kit (Qiagen 56404). The protocol was modified to include the following steps: (1) Incubated overnight with shaking in Buffer ATL with proteinase K. (2) An additional wash step with 80% ethanol prior to elution. (3) The Qiagen ATE buffer was replaced with Low TE buffer (Applied Biosystems 4389764), pre-heated to 55°C and applied to the column for ten minutes. The DNA was quantified with Quant-iT Picogreen (Invitrogen P11495).

DNA was extracted from NCI-PC35-1 and NCI-PC35-2 organoids using an AllPrep DNA/RNA Mini Kit (Qiagen 80204) according to the manufacturer’s protocol for animal cells. Qiashredder columns (Qiagen 79656) were used for the homogenization step.

### DNA whole-genome sequencing

1 μg of genomic DNA was fragmented (Covaris), end-repaired, and assembled into paired-end libraries using the Illumina TruSeq DNA Library Preparation Kit. Libraries were sequenced with 150 cycles paired-end (2 × 150) on an Illumina HiSeq 4000. Per-lane FASTQ pairs were trimmed using Trimmomatic version 0.39 and aligned to hg19 using BWA-MEM version 0.7.17. PCR duplicates were marked using the SPARK implementation of GATK MarkDuplicates version 4.1.4.1 with PICARD SetNmMdAndUqTags. Base quality score recalibration was performed using the SPARK implementation of GATK BQSRPipeline. Lane-level BAM files were merged using PICARD MergeSamFiles and GATK MarkDuplicates was run a second time with PICARD SetNmMdAndUqTags. A normal saliva sample was sequenced to a mean depth of 32.8× coverage. The tumor samples were sequenced to a mean depth of 54.5× coverage (range: 40.4× to 78.6×).

### Somatic mutation calling

MuTect2 in GATK 4.1.3.0 was used in single-sample mode to generate VCF files for each normal BAM with the disable-read-filter set to MateOnSameContigOrNoMappedMateReadFilter and max-mnp-distance set to 0. A panel-of-normals was generated using GATK GenomicsDBImport with merge-input-intervals set to true and GATK CreateSomaticPanelOfNormals. MuTect2 was next run in paired mode with each tumor sample BAM matched to its benign normal BAM from the same type of sample (FFPE or fresh) and run with the panel-of-normals (pon), filtering in real-time against mutations observed in gnomAD, and with disable-read-filter set to MateOnSameContigOrNoMappedMateReadFilter. GATK GetPileupSummaries (filtering on ExAC sites) and GATK CalculateContamination were used on each tumor BAM for filtering raw MuTect2 calls using GATK FilterMutectCalls. Finally, 8-OxoG and FFPE filtering was performed, first using GATK CollectSequencingArtifactMetrics on each tumor BAM and passing its output GATK FilterByOrientationBias with artifact-modes set to G/T and C/T. Mutations were annotated using Oncotator.

### Somatic copy number alteration calling

A joint set of copy number alterations and their clonal prevalence was determined using both GATK 4.1.3.0 and TitanCNA version 1.23.1 from whole-genome sequencing data. Using GATK, denoising was performed separately for FFPE and fresh tissues, first applying GATK CollectReadCounts for each tumor and normal BAM, and assembling a panel of normals using CreateReadCountPanelOfNormals. GATK DenoiseReadCounts was run on each tumor or normal sample using the appropriate panel of normals. GATK CollectAllelicCounts was run on each sample BAM for high-confidence 1000 Genomes Phase 1 SNP sites. Segmented copy number ratios were then calculated by using GATK ModelSegments, using denoised copy ratios for both matched tumor and normal as well as the allelic counts for each tumor sample. GATK CallCopyRatioSegments identified each region of gain or loss, per sample. TitanCNA was run using R version 3.6 on chromosomes 1-22 and X with 10kb intervals.

### Tumor phylogenetic analysis

Phylogenetic tree estimation was performed using PhyloWGS version 1.0. Prior to tree evolution, mutations input was optimized as follows: 1) MuTect2 output multi-sample VCF files were filtered to tumor-only; 2) A floating depth cutoff was applied so that mutations in a single sample must be greater than 70% of the average depth of that sample from the same patient; 3) A hard filter of 90% strand bias was imposed; 4) A combined list of all mutations for all samples from each individual were compiled with a hard filter at 10% variant allele fraction (VAF); mutations less than 10% VAF were recovered from other samples provided they were >10% VAF in at least one sample. Copy number input was optimized as follows: 1) 1-bp segments were removed from the joint output of TitanCNA and GATK; 2) high-level amplification and deep deletion events filtered from TitanCNA but present in output from the ichorCNA module were reintegrated into the .SEG file output when overlapping with GATK calls. PhyloWGS inputs per-patient were prepared using the create_phylowgs_inputs script joining each individual VCF (vardict) and CNV sample into a single set of SSM and CNV data. The corresponding SSM, CNV and parameters JSON files were then run using the multievolve script for parallel tree generation across 40 chains, using 1000 burn-in Markov chain Monte Carlo (MCMC) samples and 2500 fit MCMC iterations for a total of 100,000 potential tree structures. After tree generation, mutation and tree JSON files from the write_results script were parsed to select the tree with the lowest (most negative) log likelihood score. The best scoring tree was pruned to conservatively decrease the number of major subclones. If any given node did not have at least 5 SNVs or SSMs assigned to it, it was merged with its sibling node with the greatest number of events. If that node had no siblings, it was merged with its most immediate ancestral node, unless it was a direct descendent of the germ/normal node with no descendants, in which case it was eliminated. The subclonal composition of each node was determined by the average clonal prevalence of SSMs/CNVs assigned to each node and their relative proportion in each sequenced tumor sample.

### Immunoblots

5 x 10^5^ cells from dissociated organoids were lysed in lysis buffer (50 mM Tris (pH 8) + EDTA (10 mM) + 1% SDS) with protease and phosphatase inhibitors. Protein concentration was determined using a BCA assay (Pierce 23227). 10 μg of protein was loaded onto 4-20% Mini_PROTEAN TGX gels (Bio-Rad 456-1094) or 4-20% Mini_PROTEAN TGX Stain-Free gels (Bio-Rad 4568091). Semi-dry transfer was done with a Bio-RadTrans Blot Turbo apparatus for 30 minutes using Trans-Blot Turbo 5x Transfer Buffer (Bio-Rad10026938) except for ARID1A and ARID1B overnight - wet transfers were done. Membranes were blocked for 1 hour in 5% BSA. Overnight incubations with the primary antibodies were done at 4°C while rocking. Secondary antibody incubations were done for one hour at room temperature while rocking. Blots were developed with Clarity Western ECL Substrate (Bio-Rad 170-5061) and visualized on a Bio-Rad ChemiDoc Touch Imaging System.

### Immunofluorescent staining

Organoids were dissociated and re-plated in 2D on 16-well chamber slides (Nunc 178599) coated with 75 μg/ml poly-D-lysine (Millipore A-003-E) followed by 3% Matrigel (Corning 356231). Cells were fixed for 10 minutes in 4% formaldehyde, then rinsed three times with PBS. Cells were permeabilized and blocked for one hour in PBS/5% goat serum/0.3% Triton-X 100. The cells were then incubated in primary antibody diluted in PBS/0.5% BSA overnight at 4°C. The cells were then washed 5x fifteen minutes at room temperature in PBST and incubated with fluorochrome-conjugate secondary antibody for one hour, followed by 5x fifteen-minute washes. Coverslips were mounted with Fluoro-Gel II + DAPI (Electron Microscopy Sciences 17985-50). Slides were imaged using a Zeiss Axioscan.Z1 microscope with a plan-apochromat 20x NA 0.8 objective and a Colibri 7 LED light source. Quantification of IF images was done using the Indica Labs HALO v3.3 software running the CytoNuclear FL v2.0.12 algorithm.

### Proliferation assays

Organoids were dissociated then replated in 3D in 96 well plates. Each time-point was plated in five well replicates and incubated overnight. All time-points were then quantified at the indicated day with CellTiter Glo 3D (Promega G9682) and luminescence was measured using a Tecan infinite M200 Pro plate reader. The average fold change for each time-point relative to day-0 was calculated. Three independent experiments were performed.

### EdU-incorporation assays

Twenty-four-hour pulse: organoids were dissociated then replated in 3D overnight. The next morning 10 μM EdU (Invitrogen C10338) was added to the cultures for 24 hours. The organoids were then either immediately collected and replated in 2D for staining and imaging or they were maintained in culture for a chase period and collected at the appropriate time-point. EdU staining was performed according to the manufacturers protocol for most assays except the combination EdU/RNA-FISH assays where the following modifications were made: 1) BSA was not used in the wash buffers. 2) The incubation time in the Click-iT reaction cocktail was reduced to five minutes. Imaging was performed as describe above for immunofluorescence. Quantification of EdU was done using the Indica Labs HALO v3.3 software running either the CytoNuclear FL v2.0.12 algorithm or FISH-IF v1.2.2 algorithm. Cells were counted as EdU-positive above a minimum fluorescence value of 2,000.

Long-term incorporation assays: organoids were dissociated then replated in 3D overnight. Culture media containing 10 μM EdU was added and replaced every twenty-four hours until the organoids were collected at the appropriate time-points.

### PCA plot of AR v NE score WCM cohort

Raw FASTQ files were accessed from dbGaP phs000909.v.p1 and reanalyzed using the nextflow core RNA seq pipeline v1.0. Following the methods described in Beltran et al. (*2*), a reference AR sample was generated by using the gene expression values for genes in the AR signature from a series of three LNCaP samples sequenced at NCI/CCR. A reference neuroendocrine sample was generated by averaging the expression of neuroendocrine genes across the neuroendocrine samples from the Weill Cornell Medicine (WCM) cohort. The AR score was defined as the correlation of the expression of the sample with the AR reference sample. The integrated NEPC score is defined as the correlation between the sample and the reference neuroendocrine sample.

### scRNA-seq

Organoids growing in 3D in Matrigel and culture media in a 12-well plate were collected from the Matrigel by adding 1 mg/ml Dispase (Gibco 17105-041) to the culture for two hours and transferred to Eppendorf tubes. The organoids were pelleted by centrifuge and dissociated in 100 μl of TrypLE (Gibco 12605-028) + 100 μg/ml of DNAse-I (Sigma Aldrich DN25) for 20 minutes at 37°C with mechanical agitation every five minutes by pipette, using low retention tips. One ml of Advanced DMEM/F12 (Invitrogen 12634-02898) + 10 μM Y-27632 ROCK inhibitor (Stemcell Technologies 72307) was added to neutralize the TrypLE. The cells were then passed through a 30 μM cell strainer (Miltenyi Biotec 130-098-458) and assessed for viability and doublets before being pelleted and washed 3x in buffer (PBS + 0.04% BSA + Y-27632 (10 μM)). The cells were then counted and loaded onto the 10x Genomics Chromium platform using the 3’ v3.0 gene expression chemistry. Preparation of libraries were performed according to vendor recommendations. Single cell libraries were sequenced on either an Illumina NextSeq 500/550 instrument or an Illumina NextSeq 2000 instrument. Data was processed using the 10x Genomics cellranger pipeline to demultiplex reads and then align those reads to the GRCh38 reference genome. Gene barcode matricies were generated using the cellranger pipeline from 10x Genomics aligned against grch38. An in-house single cell processing pipeline was used to standardize analysis across all samples which follows the methodology laid out in the Bioconductor single cell analysis book(*35*). Briefly, gene barcode matrices were read into R and doublets were detected and removed using scDblFinder. Additional quality control was applied using the scran and scatter packages, using the addPerCellQC function and filtering out cells that were identified as outliers using the Outlier function for mitochondrial gene content, lower number of reads, and lower number of detected genes. Initial dimensional reduction was performed using GLMPCA from the scry package on all genes(*36*). UMAPs were generated from three independent experiments for PC35-1 and PC35-2 and two experiments for LuCaP 145.2. Mutual nearest neighbor correction was performed to correct for batch effects on the principal components, and the corrected top 30 principal components were used to generate the UMAP(*37*). For PC44, UMAP was performed on the top 30 principal components from one experiment. Monocle3’s graph-based clustering using leiden community detection with a q value cutoff of 0.05 was used to identify clusters and larger partitions. Marker gene detection was performed using the score markers function from scater. Cell cycle state was inferred using cyclone.

### scRNA-seq signature scores

Signature scores for individual cells were generated by running PCA on batch corrected and normalized expression values from all single cell RNA sequencing samples using only the genes in published signatures. Signature composition and references are available in supplemental table X.. The signature value is the loading for a particular cell from the first principal component.

### RNA velocity

RNA velocity was calculated independently on each sample using the default settings in velocyto as described in the orchestrating single cell analsyis with bioconductor(*35*). RNA velocity vectors were generated using batch corrected principal components to embed on the UMAP.

### CellTag analysis

Organoids were collected and dissociated to single cells for transduction with a lentiviral library of CellTags. The CellTag library (CTL) was prepared according to Biddy et al. (*18*). Lentivirus was made by transfecting Lenti-X 293T cells (Clontech 632180) with CTL plasmids plus psPAX2 and VSV-G packaging plasmids using Lipofectamine 2000 (Invitrogen 11668019). The transfection mix was applied to the cells for six hours then removed and replaced with lentiviral collection media: DMEM + 10% FBS(HyClone) + 1.1% BSA + HEPES (10 mM) + sodium pyruvate (10 mM) + Primocin (Invivogen ant-pm-1). The lentivirus was collected in two batches at 48 and 72 hours and pooled together, then spun for 5 minutes at 1000 x g to pellet debris. The supernatant was then passed through a 0.45 μM PES membrane filter. The lentivirus was concentrated 100-fold by ultracentrifuge: four hours at 4°C at 20,000 x g with low acceleration and then resuspended in PBS, aliquoted and stored at -80°C. For transduction, 5 x 10^5^ cells were combined with 3.5 μl of lentivirus and 2 μl of LentiBOOST (Sirion Biotech) in 2 ml of culture media. The cells/lentivirus were transferred to one well of a 6-well plate coated with 3% Matrigel and centrifuged at 1,000 x g, low acceleration, for 90 minutes at 32°C. The plate was then incubated overnight at 37°C. The next morning the cells were detached from the plate with TrypLE, collected and counted, then re-plated in 3D in multiple wells of a 24-well plate at different concentrations ranging from 5 x 10^3^ – 1 x 10^5^ in order to maximize recovery of the targeted 10,000 - 15,000 cells desired for loading onto the 10x Genomics platform. The cells were kept in culture for four weeks, changing the media twice/week. Each well of organoids was then collected and processed as described above in the “scRNA-seq” paragraph of this methods section. After counting, we determined that 5 x 10^4^ cells/well yielded the ideal 15,000 cells after processing. 15,000 single cells were loaded onto the 10x Genomics Chromium platform as described above. Single cell libraries were sequenced as described above. Raw single cell FASTQs were aligned to a custom reference including the EGFP construct used in the vector for the cell tags (*18*). Reads were filtered to include only sequences that aligned to EGFP. The CellTagR package was used with barcode correction relying on starcode to call clones(*38*). Cells were considered clones if they shared at least two celltags and their jaccard similarity exceeded 0.7 as specified in the documentation. To project clones onto UMAP embeddings, segments were drawn between cells that were called clones.

### RNA-FISH

Organoids were dissociated and 75,000 cells were replated overnight in 2D on 12 mm round #1 coverglass (Electron Microscopy Sciences 72231-01) coated with 75 μg/ml Poly-D-lysine followed by 3% Matrigel. The cells were washed in PBS, fixed in 4% formaldehyde for 10 minutes at room temperature, and finally washed twice in PBS. The cells were permeabilized in 70% ethanol for at least one hour at 4°C, the ethanol was removed, and Wash Buffer A (Biosearch Technologies SMF-WA1-60) was added and incubated at room temperature for five minutes. For staining, the Stellaris RNA-FISH probes, diluted in Hybridization Buffer (Biosearch Technologies SMF-HB1-10) plus 10% Deionized Formamide (Millipore 4610), were added to the cells and incubated overnight in a humidified chamber at 37°C. The cells were washed in Wash Buffer A for 30 minutes at 37°C in the dark, then counter-stained with 5 ng/ml DAPI diluted in Wash Buffer A in the dark at 37°C for 30 minutes. The cells were washed in Wash Buffer B (Biosearch Technologies SMF-WB1-20) for 5 minutes at room temperature in the dark, the cover glass was mounted onto a slide with ProLong Gold antifade reagent (Invitrogen P36934), allowed to dry and stored at -20°C in the dark. For RNA-FISH/EdU and RNA-FISH/IF combined assays, the RNA-FISH hybridization was done first up to/including the Wash Buffer B step. The cells were rinsed twice in PBS and stained for EdU or stained with antibodies for IF. For EdU incorporation/staining, see the above methods section for details. For IF, the blocking step was excluded, and antibodies were diluted in PBS. Imaging was done with a Nikon Ti2 microscope equipped with a CFI Plan-Apochromat 60x NA 1.4 oil immersion objective, Lumencor Sola SE 365 FISH light engine, and Photometrics Prime BSI sCMOS camera. A maximum intensity projection was created from a 3.6 μM 13 step Z stack for each field of view. Quantification of RNA-FISH and combined assay images was done using Indica Labs HALO v3.3 software running the FISH-IF v1.2.2 algorithm. For RNA-FISH scatter plots, total FISH counts were plotted. For the RNA-FISH/IF combined assays, total FISH counts were plotted against raw IF intensity values. For the RNA-FISH/EdU drug-treated assays, a minimum threshold of five spots (transcripts) per cell was set to call a cell positive for a given FISH marker. Cells were counted as EdU-positive above a minimum fluorescence value of 2,000. The EZH2 probe set was ordered from the Stellaris Design Ready Probe Sets (Biosearch Technologies VSMF-2123-5). All other RNA-FISH probe sets were custom designed using the Stellaris Probe Designer tool at the biosearchtech.com website, and QC’d for specificity using the UCSC genome browser BLAT function. The custom designed RNA-FISH probe-set sequence information is in Table S1.

### CopyKAT

Copy number variation was computed using CopyKat (*19*) with default setting and *cell.line* mode enabled. Briefly, raw counts from scRNAseq experiments were used as input to CopyKat. CopyKat clusters were generated by unsupervised hierarchical clustering of the CNV results using the function *hclust* with ward.D linkage function on the cell distance matrix computed using the *dist* function calculated with “euclidean” method. Copykat clusters were assigned based on the number of UMAP clusters using *cutree* function.

### scATAC-seq

Organoids were collected and dissociated as described above in the “scRNA-seq” paragraph of this methods section. Single cell suspensions of 2.5 x 10^5^ cells were spun down and resuspended in 100 μl of cold ATAC lysis buffer (10 mM Tris(pH 7.4) + 10 mM NaCl + 3 mM MgCl2 + 1% BSA + 0.1% Tween-20), pipetted up/down 10x, incubated on ice for five minutes, and finally pipetted an additional 5x before adding 1 ml of ATAC Wash Buffer (10 mM Tris(pH 7.4) + 10 mM NaCl + 3 mM MgCl2 + 1% BSA + 0.1% Tween-20 + 0.1% NP40 + 0.01% digitonin). The cells were then pelleted at 500 x g for 5 minutes at 4°C. All of the wash buffer was removed, and the nuclei were resuspended in 50 μl of Nuclei Buffer (10x Genomics PN-2000153/2000207). Single nuclei suspensions were transposed before being partitioned on the 10x Genomics Chromium platform using the Single Cell ATAC v1.1 chemistry (10x Genomics). Preparation of libraries were performed according to vendor recommendations.

Single cell atac sequencing was processed using the cellranger scatac pipeline from 10x Genomics. Additional analysis was performed using the ArchR library using 250,000 features for the latent semantic indexing(*39*). Inferred transcription factor activity was generated using the method included in ArchR for generating ChromVAR deviations Z scores. The score markers function was applied and performs a competitive ranking of features using three statistical tests; Welch’s t test, Wilcoxon rank sum test, and a binominal test. Features that were ranked highly in all three tests were considered. Assignment of clusters to phenotypes was performed using the differential accessibility metric as described(*24*) as an input into GSEA(*40*) with references being comprised of the transcription factors identified as being regulators of prostate cancer phenotypes(*22, 24, 25*).

### Dose-response assays

Organoids were dissociated then replated in 3D at 3,000 cells/well in 384 well plates. Drugs were prepared by two-fold serial dilutions starting at 10 μM and spanning 11 concentrations, plus an additional vehicle control. All treatments were done in replicates of five. The cells were treated twice per week for two weeks, then quantified with CellTiter Glo 3D and luminescence was measured using a Tecan infinite M200 Pro plate reader. Data is shown as an average of three independent experiments.

### Xenograft tumor study

The animal study was performed according to the protocol approved by the NCI-Bethesda Animal Care and Use Committee. The organoid-derived xenograft (ODX) model was established initially from NCI-PC35-1 organoids subcutaneously injected in NOD scid gamma (NSG) mice, and subsequently maintained by serial passage of tumor fragments in NSG mice. For the experiment, 2 mm tumor fragments were implanted subcutaneously in NSG mice. When the tumors reached an average size of 0.3 cm^3^ the mice were randomized into four treatment groups of five mice/group. Mice in the castrated groups were castrated by orchiectomy concurrent with the start of drug treatment. Mice were drugged once daily, five days/week by oral gavage with 30 mg/kg of alisertib (Fig. 7) suspended in vehicle (10% 2-hydroxypropyl-*β*-cyclodextrin, 1% sodium bicarbonate in water) or twice daily with 20 mg/kg alisertib (Fig. S10). Mice in the vehicle control group were treated on the same schedule. Tumor volumes were measured twice/week. The study was terminated after nine weeks when the control group reached the maximum allowable burden of 2 cm^3^. Tumors were harvested and fixed in 4% formaldehyde overnight then transferred to 70% ethanol.

### Antibodies

| Target | Company | Catalog number | Assay |
| --- | --- | --- | --- |
| AR | Abcam | ab133273 | IF/Western |
| AR | Cell Signaling | 5153 | IHC |
| ARID1A | Cell Signaling | 12354 | Western |
| ARID1B | Abcam | ab57461 | Western |
| BrdU | Abcam | ab6326 | IHC |
| CHGA | Invitrogen | MA5-13096 | IF/IHC |
| EZH2 | Cell Signaling | 5246 | Western |
| EZH2 phospho-S21 | Bethyl Laboratories | IHC-00388 | Western |
| EZH2 phospho-T345 | Active Motif | 61242 | Western |
| EZH2 phospho-T350 | Gift from Amina Zoubeidi | NA | Western |
| p27 | BD Biosciences | 610241 | Western |
| p53 | Cell Signaling | 2524 | Western |
| PTEN | Cell Signaling | 9188 | Western |
| RB | Cell Signaling | 9309 | Western |
| SYP | Agilent | M731529-2 | IHC |

## Data availability

The sequence information for all RNA-FISH probe sets is located in Supplementary Information Table 1. The WGS, scRNA-seq, and scATAC-seq data have been deposited in GEO.

