## Supplemental Figures for "The origin and dynamics of cellular heterogeneity vary across lineage subtypes of castrate resistant prostate cancer"

A

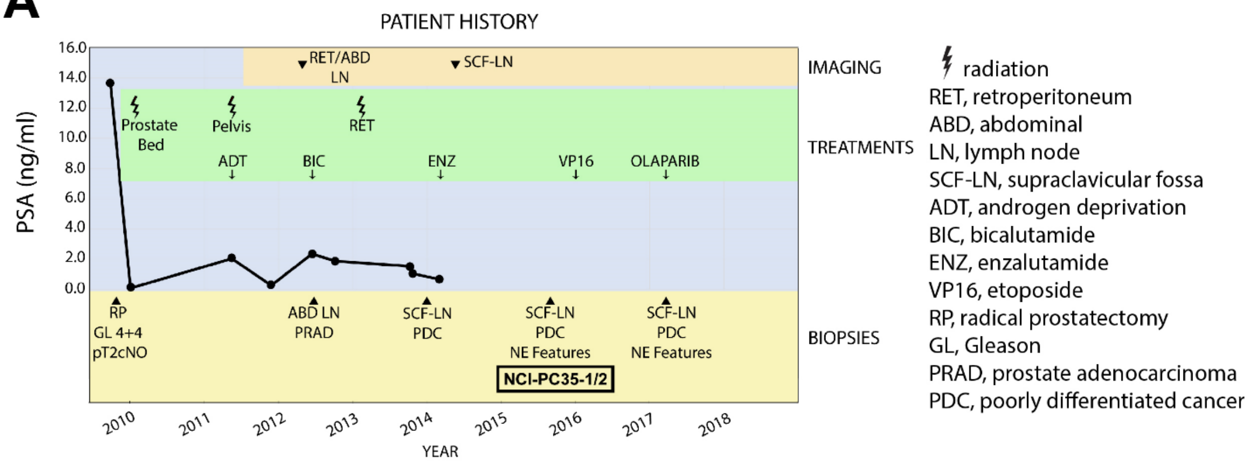


| Primary tumor | prostate |
|---|---|
| Histologic type: | Adenocarcinoma |
| Gleason Score: | 4 + 4 = 8 |
| Tumor Quantification : | Tumor involves greater than 50% of both the right and left lobes |
| Margins: | Uninvolved |
| Extraprostatic extension: | Absent |
| Seminal vesicle invasion : | Absent |
| Number of regional lymph nodes examined: | 3 |
| Number of lymph nodes with metastasis: | 0 |

| Site of Metastasis | subclavicular LN |
|---|---|
| Dx | Metastatic poorly differentiated carcinoma with NE features. |
| Hx | goserelin, leuprolide, bicalutamide, enzalutamide |
| Markers | PAP, CDX2, chromogranin , synaptophysin, PSA(rare/faint) |

B

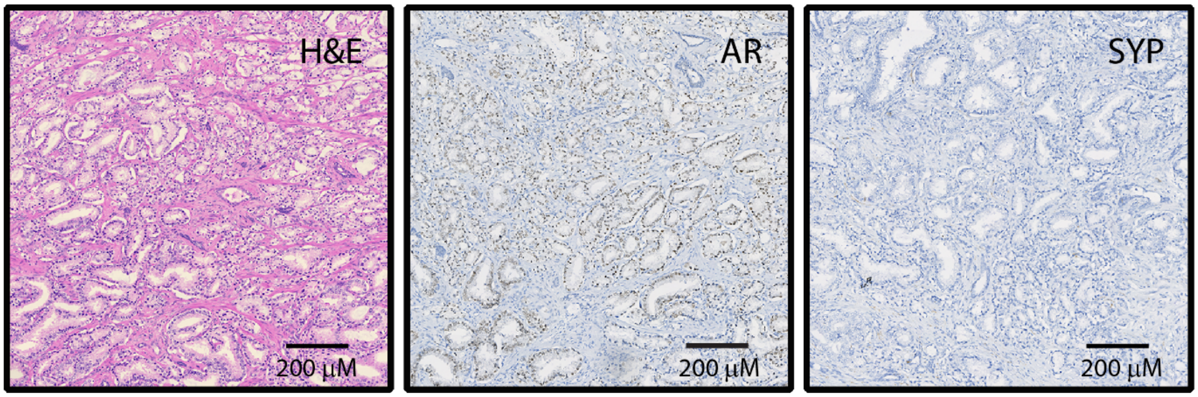

**Figure S1. Patient treatment history. Related to Figure 1.** (A) Indicates the course of treatment and diagnosis of the patient. The time of biopsy collection from which the NCI-PC35 organoid lines were established is indicated. (B) Primary prostate tumor sections stained with H&E or antibodies against AR or the neuroendocrine marker synaptophysin (SYP).

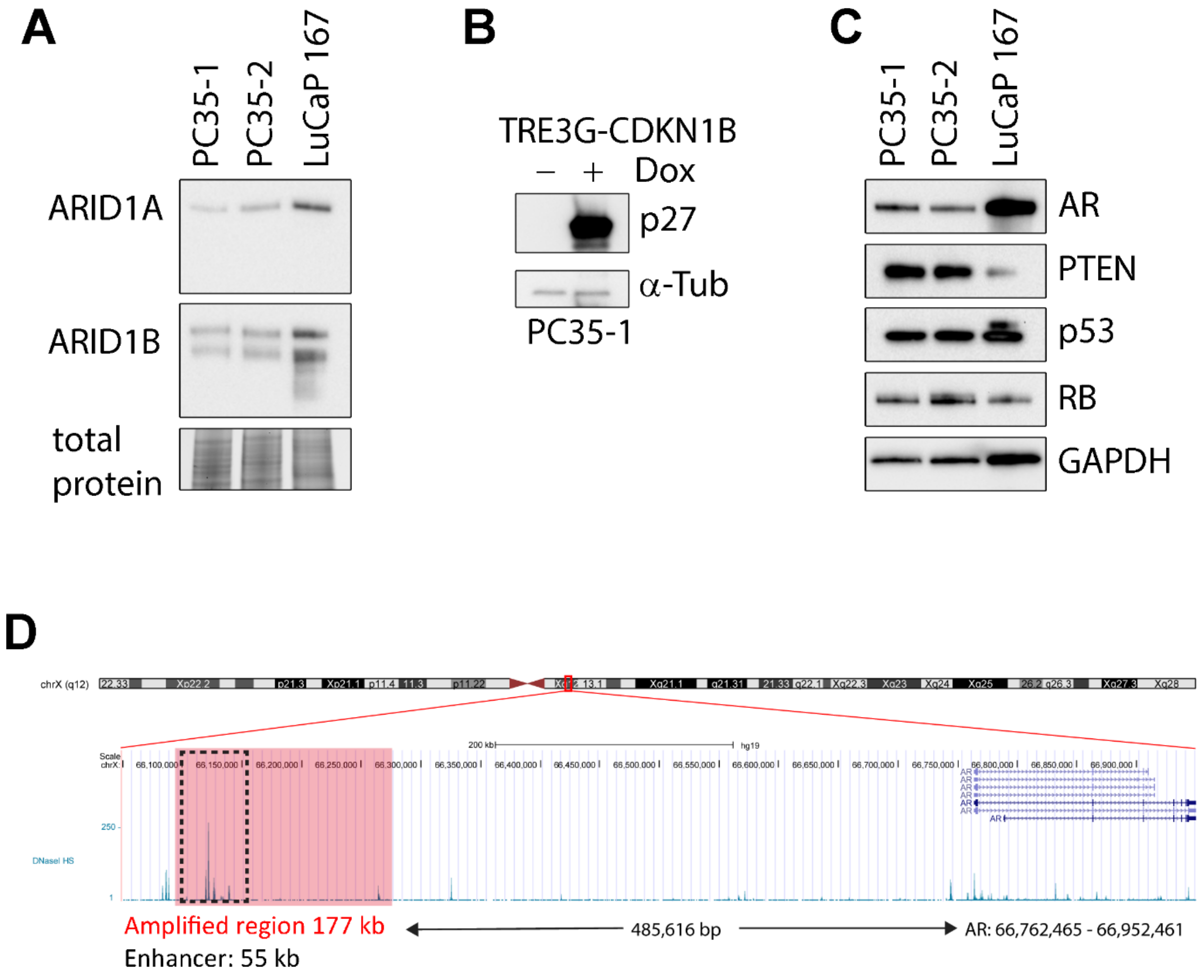


E

***ARID1A* and *ARID1B* mutations**

*Breakdown of mutational status from public data*

| Tumor Source | Wild Type | Alternative | Single Copy Loss |
|---|---|---|---|
| | Number of cases | | |
| *ARID1A* | | | |
| Primary | 639 (93.3%) | 9 (1.31%) | 37 (5.4%) |
| Metastatsis | 364 (78.8%) | 28 (6.06%) | 70 (15.2%) |
| *ARID1B* | | | |
| Primary | 624 (90.7%) | 14 (2.03%) | 50 (7.27%) |
| Metastatsis | 321 (69.2%) | 70 (15.1%) | 73 (15.7%) |

Data compiled from prostate cancer studies in cBioPortal and WCDT

**Figure S2. Genetic alterations of PC35 organoids affect protein levels. Related to Figure 1.** (A) Western blots showing protein expression of ARID1A and ARID1B from lysates of PC35-1 and PC35-2 relative to WT LuCaP 167 control. Total protein is shown as a loading control. (B) Western blots showing p27 protein expression in PC35-1. Endogenous expression (-Dox) compared to expression from a positive control, doxycycline-inducible expression vector (+Dox). $\alpha$-tubulin loading control. (C) Western blots showing protein expression of AR, PTEN, p53, and RB in PC35-1, PC35-2 and LuCaP 167 organoids. GAPDH loading control. (D) Region of tandem duplication that encompasses the AR enhancer. (E) Number of cases from prostate cancer studies in cBioPortal and the WCDT with variants in *ARID1A* and *ARID1B*. Alternative refers to mutations predicted to alter protein coding function.

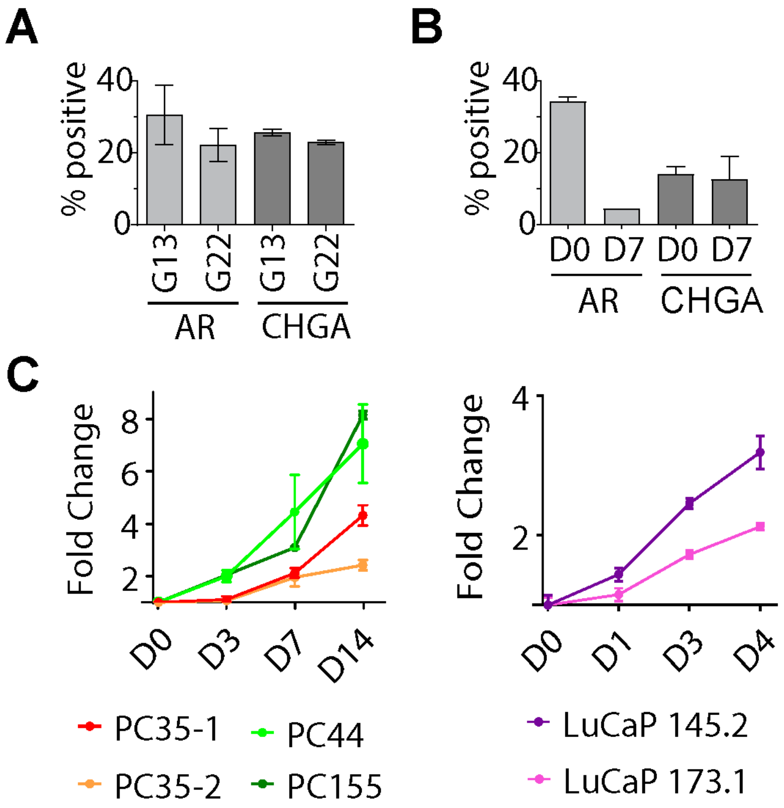
A
% positive
40
20
0
G13
G22
G13
G22
AR
CHGA
B
% positive
40
20
0
D0
D7
D0
D7
AR
CHGA
C
Fold Change
8
6
4
2
D0
D3
D7
D14
PC35-1
PC44
PC35-2
PC155
Fold Change
4
2
D0
D1
D3
D4
LuCaP 145.2
LuCaP 173.1

**Figure S3. Growth kinetics of organoid models and of specified PC35 lineage populations. Related to Figure 1.** (A) IF staining of CT35-1 organoids with antibodies against the indicated proteins. Generation 13 and 22 organoids were dissociated and quantified as single cells then plotted as percent EdU-positive of total marker-positive cells. (B) IF combined with 5-ethynyl-2'-deoxyuridine (EdU) pulse-chase assay of PC35-2. Organoids were pulsed with EdU (10μM) for 24 hours (D0) and chased for seven days (D7). The organoids were dissociated and quantified as single cells and plotted as percent-positive of total cells. (C) Relative fold change in growth over time (D = day) for the indicated organoid lines, quantified by CellTiter Glo 3D.

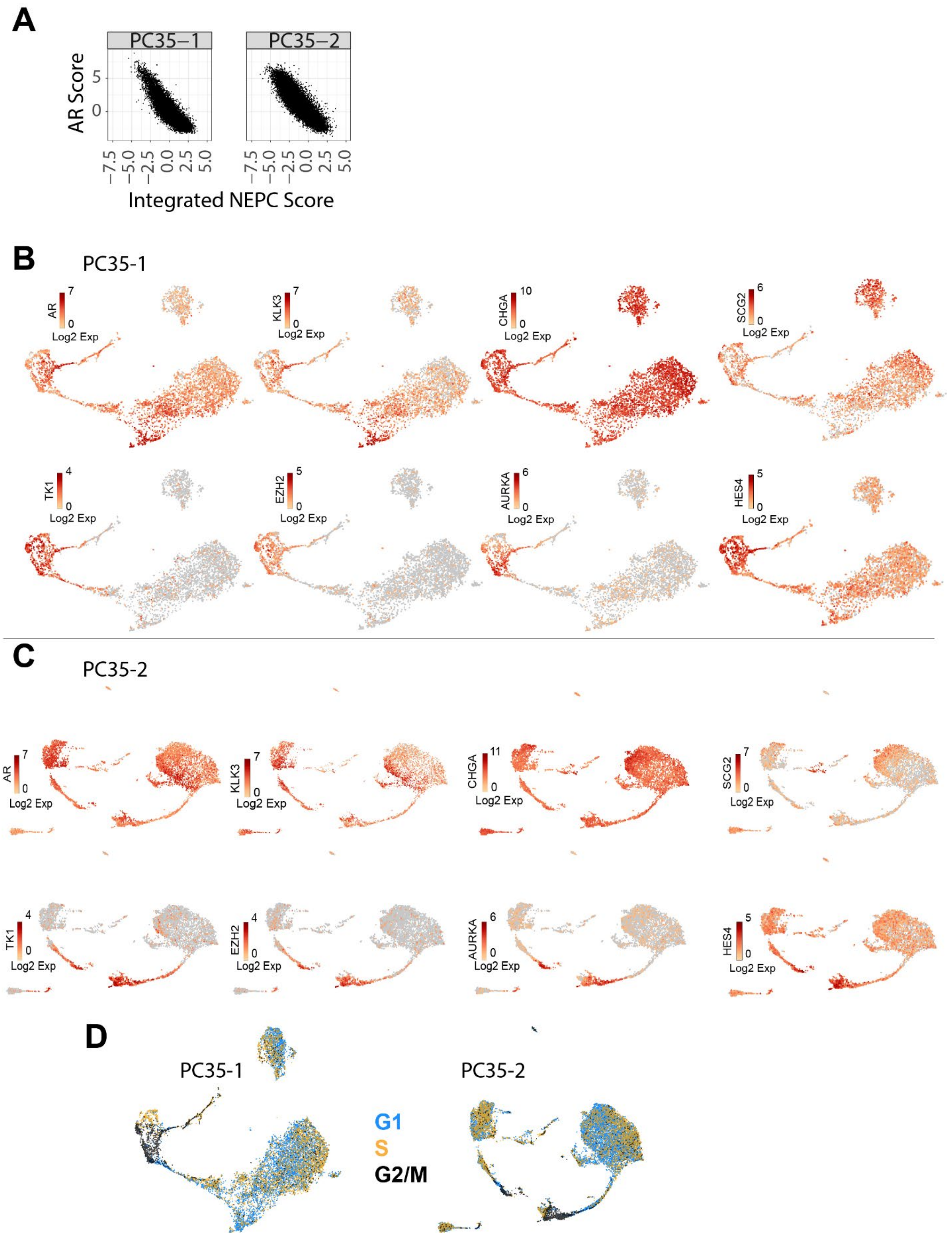
A
PC35−1
PC35−2
AR Score
5
0
−7.5 −5.0 −2.5 0.0 2.5 5.0
Integrated NEPC Score
B
PC35-1
AR
KLK3
CHGA
SCG2
TK1
EZH2
AURKA
HES4
Log2 Exp
C
PC35-2
AR
KLK3
CHGA
SCG2
TK1
EZH2
AURKA
HES4
Log2 Exp
D
PC35-1
PC35-2
G1
S
G2/M

**Figure S4. Single-cell transcriptomic analysis of organoid models. Related to Figure 2.** (A) Each cell from PC35-1 and PC35-2 scRNA-seq transcript data is plotted by neuroendocrine score (x axis) and AR score (y axis). (B) PC35-1 and (C) PC35-2, Log2 expression of the indicated lineage markers overlaid on the UMAPs. ARPC (AR, KLK3); NEPC (CHGA, SCG2); proliferation/stem markers (TK1, EZH2, AURKA, HES4) (D) Cell cycle phase transcriptional signature overlaid on UMAPs of PC35-1 and PC35-2.

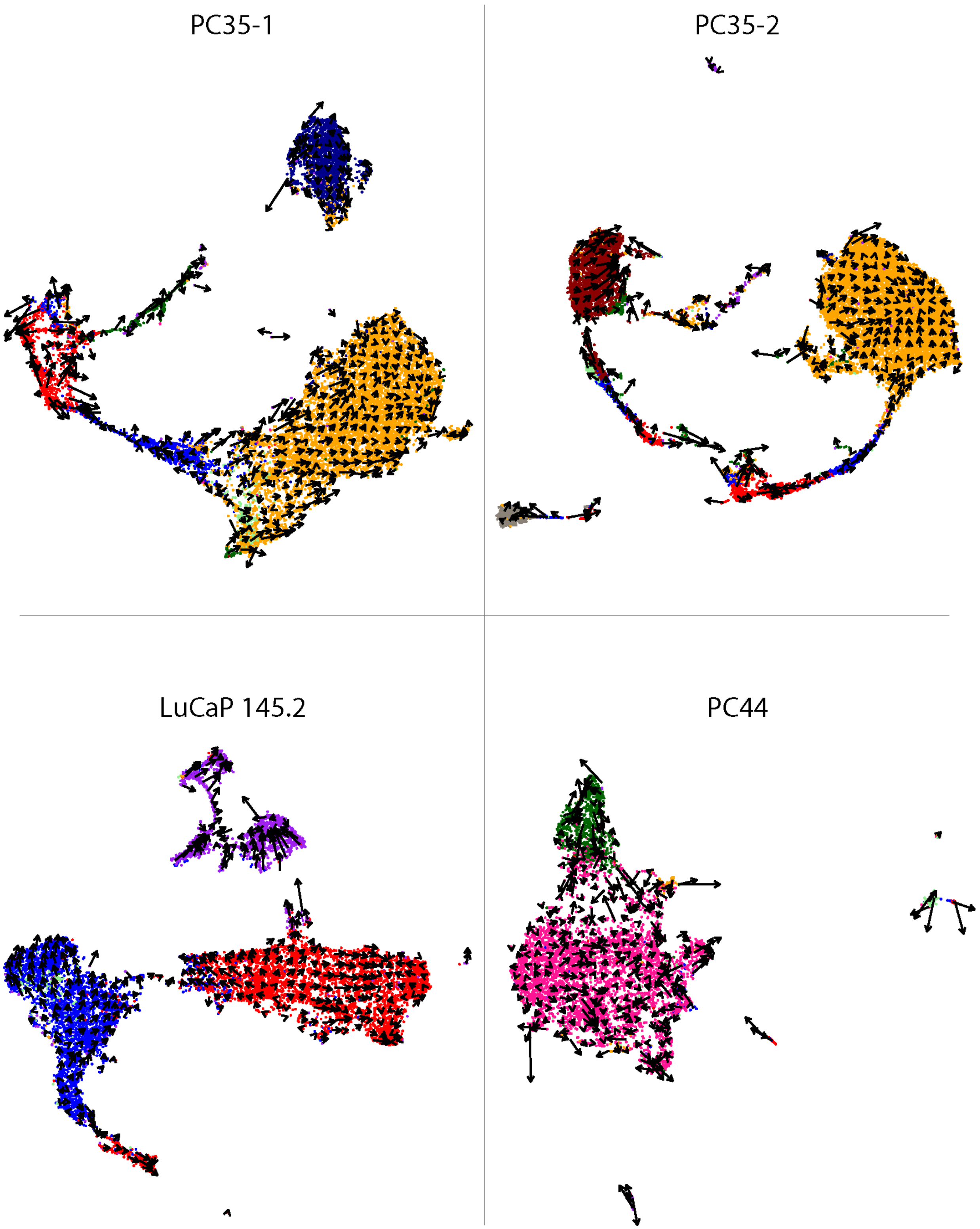
PC35-1
PC35-2
LuCaP 145.2
PC44

**Figure S5. RNA-velocity analysis identifies subpopulations in the PC35 organoid models with features of stem/progenitor populations. Related to Figure 2.** RNA velocity vectors are embedded onto the UMAPs for each model. The direction of each velocity points toward the future state of locally averaged vector fields. The length indicates the magnitude of difference between states.

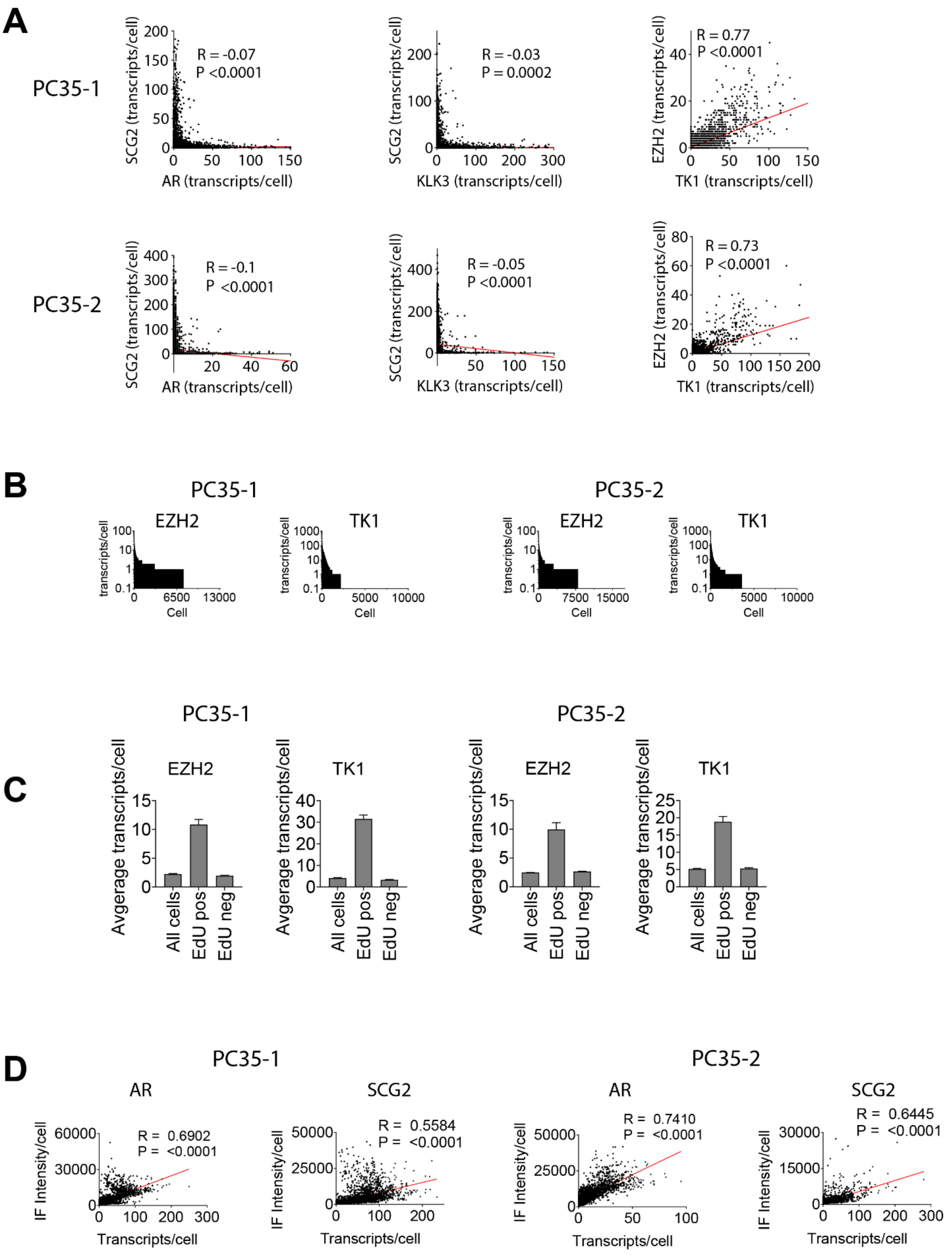
A
PC35-1
R = -0.07
P <0.0001
SCG2 (transcripts/cell)
AR (transcripts/cell)
R = -0.03
P = 0.0002
KLK3 (transcripts/cell)
R = 0.77
P <0.0001
EZH2 (transcripts/cell)
TK1 (transcripts/cell)
PC35-2
R = -0.1
P <0.0001
R = -0.05
P <0.0001
R = 0.73
P <0.0001
B
PC35-1
EZH2
TK1
PC35-2
transcripts/cell
Cell
C
PC35-1
PC35-2
EZH2
TK1
Avgerage transcripts/cell
All cells
EdU pos
EdU neg
D
PC35-1
PC35-2
AR
SCG2
R = 0.6902
P = <0.0001
R = 0.5584
P = <0.0001
R = 0.7410
P = <0.0001
R = 0.6445
P = <0.0001
IF Intensity/cell
Transcripts/cell

**Figure S6. Single-molecule RNA-FISH analysis of PC35-1 and PC35-2 corroborates the results of scRNA-seq and pathological analyses. Related to Figure 2.** (A) Scatter plots of marker genes expression in individual cells. Each point represents a single cell plotted as the number of transcripts per cell for the genes indicated on X and Y. (B) Distribution plots of EZH2 and TK1 transcripts per cell quantified by RNA-FISH. Each cell is plotted on the X axis and the number of transcripts is shown on the Y axis in log scale. (C) Quantification of marker gene expression in dividing and non-dividing cells. PC35-1 and PC35-2 organoids were pulsed with EdU for 24 hours, dissociated and stained for EdU incorporation and transcript abundance. The average number of transcripts per cell for the indicated genes are shown for three populations of cells with respect to EdU-incorporation status. (D) Scatter plots of combined RNA-FISH/IF assays for PC35-1 and PC35-2. The number of transcripts per cell for the indicated gene is plotted against the immuno-fluorescence intensity per cell for the protein product of that gene (see Methods). For all scatter plots, the Pearson correlation coefficient was calculated and shown as R. The line was fit by simple linear regression. All bar plots are shown as the mean of independent experiments ± SEM.

| Celltag Tabulation | | | | | | |
|---|---|---|---|---|---|---|
| | | Self Renewing | | Transitioning | | |
| | Cluster | Count | Percent | Count | Percent | Number of cells in cluster |
| PC35-1 | | | | | | |
| | 1 | 9 | 0.80% | 18 | 1.59% | 1130 |
| | 2 | 11 | 1.02% | 16 | 1.48% | 1083 |
| | 3 | 6 | 0.07% | 11 | 0.13% | 8288 |
| | 5 | 0 | 0.00% | 5 | 1.89% | 264 |
| | 6 | 0 | 0.00% | 3 | 0.57% | 528 |
| | 8 | 2 | 0.10% | 0 | 0.00% | 2030 |
| | 9 | 0 | 0.00% | 2 | 5.00% | 40 |
| Total | — | 28 | — | 55 | — | 13,363 |
| Percent | — | 0.21% | — | 0.41% | — | 100.00% |
| PC35-2 | | | | | | |
| | 1 | 36 | 2.70% | 25 | 1.88% | 1333 |
| | 2 | 22 | 1.80% | 27 | 2.21% | 1224 |
| | 3 | 2 | 0.02% | 4 | 0.04% | 10664 |
| | 5 | 0 | 0.00% | 3 | 1.60% | 188 |
| | 7 | 7 | 0.22% | 5 | 0.16% | 3118 |
| | 9 | 0 | 0.00% | 1 | 7.69% | 13 |
| | 10 | 0 | 0.00% | 1 | 0.12% | 822 |
| Total | — | 67 | — | 66 | — | 17,362 |
| Percent | — | 0.39% | — | 0.38% | — | 100.00% |
| LuCaP 145.2 | | | | | | |
| | 1 | 78 | 1.51% | 122 | 2.37% | 5155 |
| | 2 | 39 | 1.03% | 90 | 2.37% | 3793 |
| | 4 | 1 | 0.05% | 0 | 0.00% | 2137 |
| | 5 | 0 | 0.00% | 4 | 2.35% | 170 |
| | 8 | 0 | 0.00% | 1 | 16.67% | 6 |
| Total | — | 118 | — | 217 | — | 11,261 |
| Percent | — | 1.05% | — | 1.93% | — | 100.00% |
| PC44 | | | | | | |
| | 1 | 2 | 0.86% | 4 | 1.72% | 233 |
| | 2 | 1 | 0.65% | 5 | 3.23% | 155 |
| | 5 | 0 | 0.00% | 3 | 1.49% | 202 |
| | 9 | 117 | 2.30% | 11 | 0.22% | 5094 |
| Total | — | 120 | — | 23 | — | 5,684 |
| Percent | — | 2.11% | — | 0.40% | — | 100.00% |

**Figure S7. Tabulation of cells with CellTags detected in each model and cluster**. **Related to Figure 3.** Number and percentage of cells with CellTags within each cluster are reported. Cells with CellTags are identified as Self Renewing or Transitioning populations. A cell was considered Self Renewing if all cells with a specified CellTag were found in the same cluster, and Transitioning if cells with a specified CellTag were found in multiple clusters.

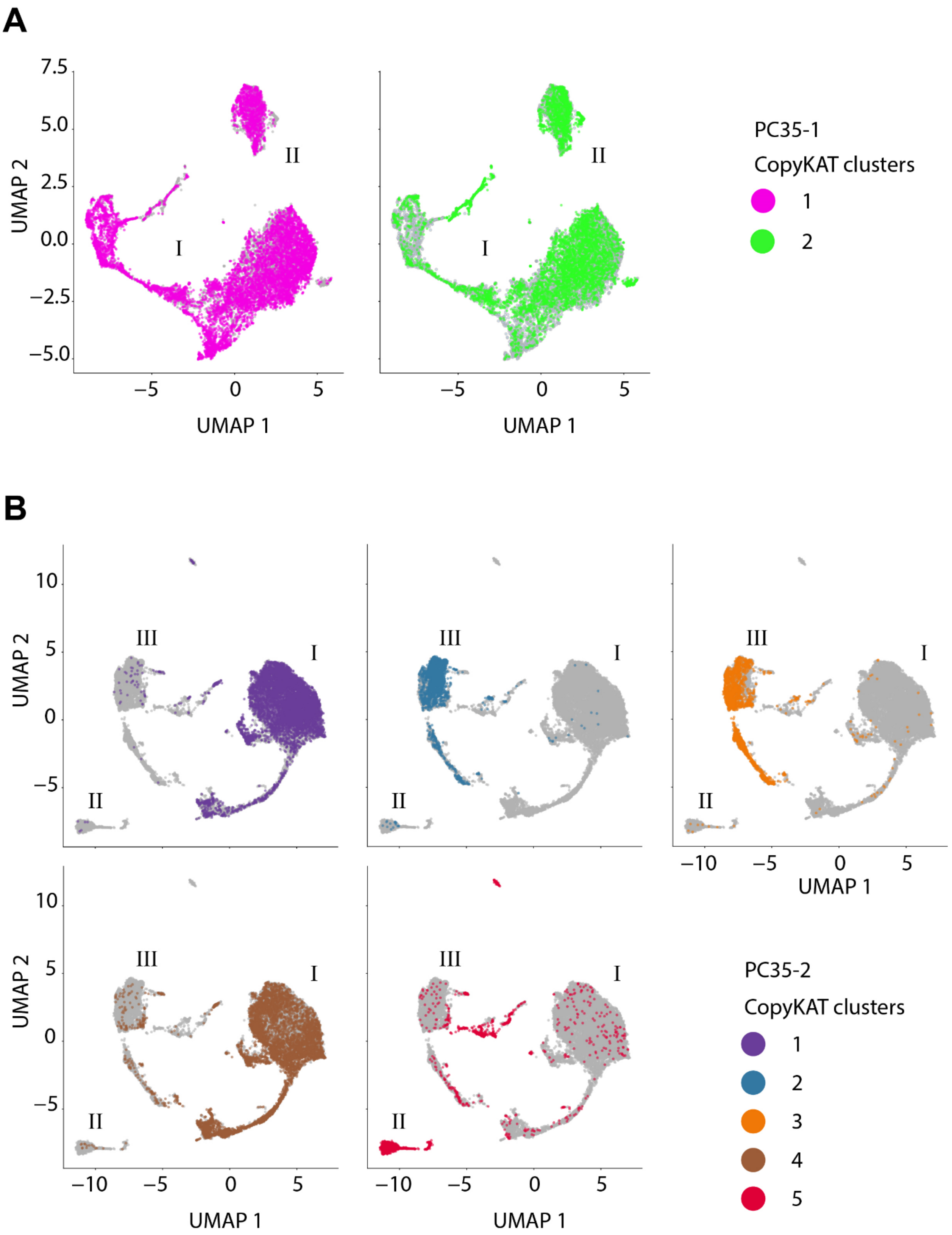

A
7.5
5.0
2.5
0.0
−2.5
−5.0
UMAP 2
II
I
−5
0
5
UMAP 1
PC35-1
CopyKAT clusters
1
2
B
10
5
0
−5
III
I
II
−10
−5
0
5
UMAP 1
UMAP 2
PC35-2
CopyKAT clusters
1
2
3
4
5

**Figure S8. PC35 major clusters defined by transcriptional profile (phenotype) are partially linked to genotype, while genotypes are not biased to specific regions within UMAP clusters. Related to Figure 3.** (A) PC35-1 and (B) PC35-2 UMAPs showing major clusters (I, II, III). Each cell of the genomic CNV-determined subclonal populations (CopyKAT clusters) is indicated by color on a UMAP shown for each CopyKAT cluster.

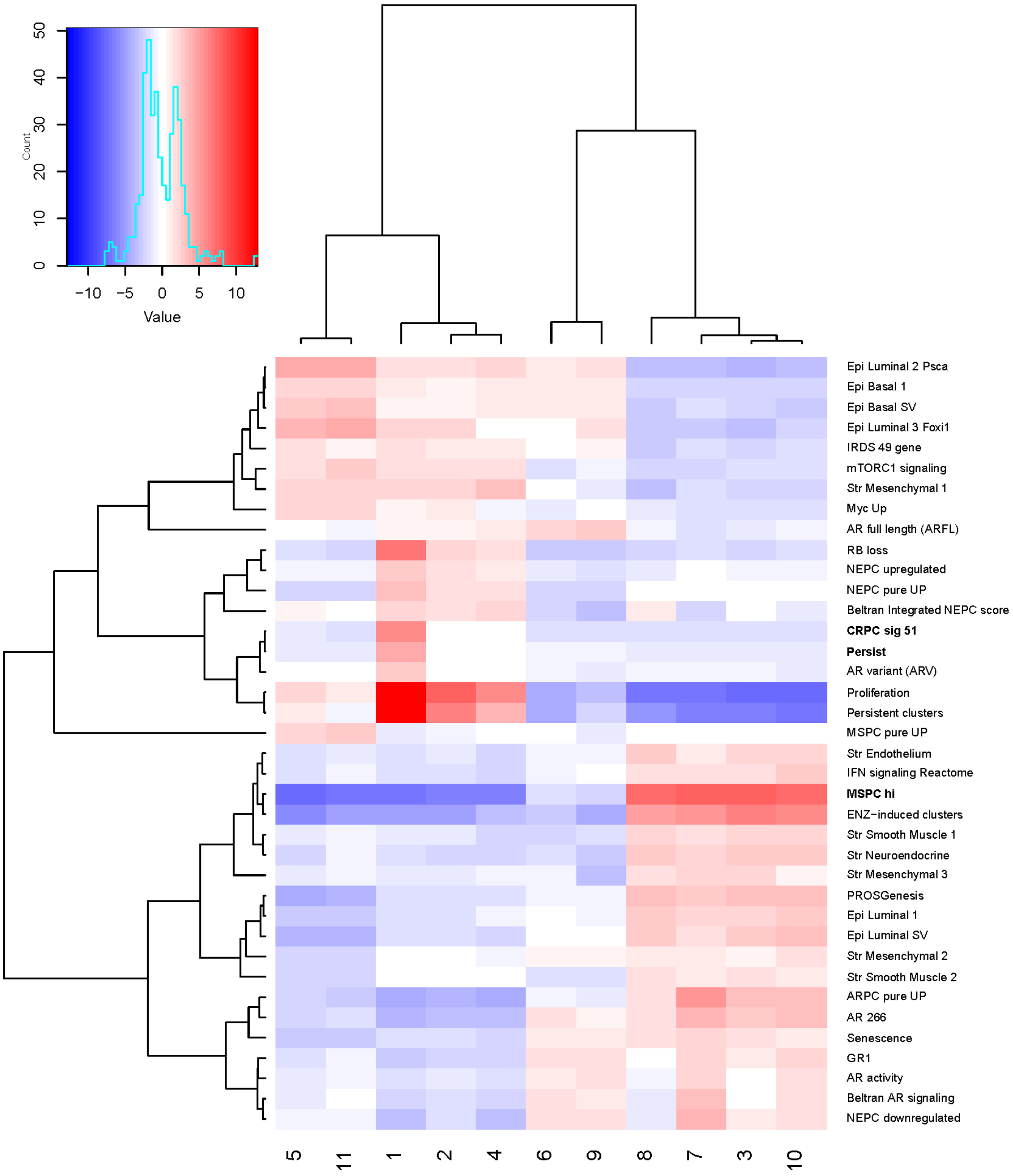

Count
Value
Epi Luminal 2 Psca
Epi Basal 1
Epi Basal SV
Epi Luminal 3 Foxi1
IRDS 49 gene
mTORC1 signaling
Str Mesenchymal 1
Myc Up
AR full length (ARFL)
RB loss
NEPC upregulated
NEPC pure UP
Beltran Integrated NEPC score
CRPC sig 51
Persist
AR variant (ARV)
Proliferation
Persistent clusters
MSPC pure UP
Str Endothelium
IFN signaling Reactome
MSPC hi
ENZ−induced clusters
Str Smooth Muscle 1
Str Neuroendocrine
Str Mesenchymal 3
PROSGenesis
Epi Luminal 1
Epi Luminal SV
Str Mesenchymal 2
Str Smooth Muscle 2
ARPC pure UP
AR 266
Senescence
GR1
AR activity
Beltran AR signaling
NEPC downregulated
5
11
1
2
4
6
9
8
7
3
10

**Figure S9. Cluster Phenotypes. Related to Figure 4.** Heat map showing mean signature scores for each cluster across all samples. Samples and scores were clustered using correlation distance.

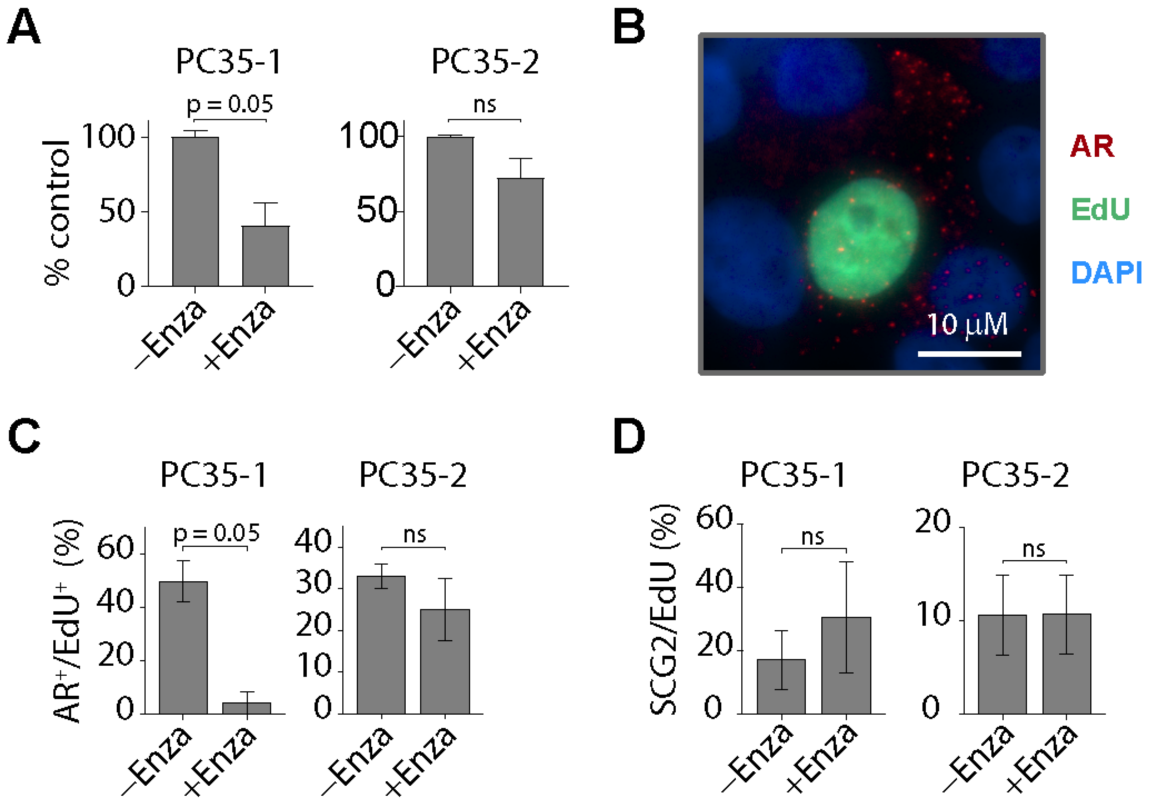
A
PC35-1
PC35-2
p = 0.05
ns
% control
100
50
0
–Enza
+Enza
B
10 μM
AR
EdU
DAPI
C
AR+/EdU+ (%)
60
40
20
30
10
D
SCG2/EdU (%)

**Figure S10 related to Figure 6. PC35 organoids show subpopulation-specific sensitivity to AR inhibition** (A) PC35-1 and PC35-2 organoids treated for six weeks with enzalutamide (10 μM). Relative cell numbers were quantified with CellTiter Glo 3D and plotted relative to the control. (B) A representative image of a combination RNA-FISH/EdU assay on PC35-1 organoids. The organoids were pulsed for 24 hours with EdU prior to collection and then dissociated and replated in 2D on cover slips and stained for AR and EdU. (C) PC35-1 and PC35-2 organoids were treated for six weeks with enzalutamide (10 μM). Organoids were pulsed with 10 μM EdU for 24 hours prior to collection, then dissociated and replated in 2D on cover slips and stained for AR expression by RNA-FISH. EdU incorporation status (positive or negative) was determined for each cell (see Methods). The data was plotted as the percentage of EdU-positive cells that also expressed AR in each treatment condition. (D) PC35-1 and PC35-2 organoids were treated as in (C) and stained for SCG2 expression by RNA-FISH. EdU incorporation status (positive or negative) was determined for each cell. The data was plotted as the percentage of EdU-positive cells that also expressed SCG2 in each treatment condition.

**A**

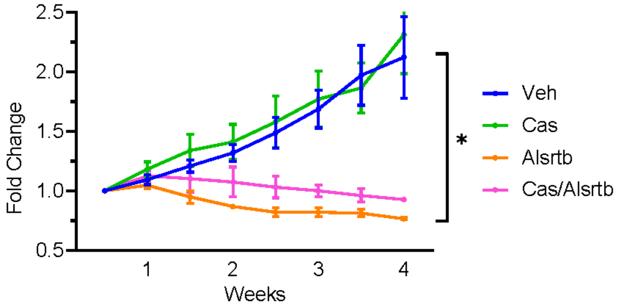
2.5
2.0
1.5
1.0
0.5
Fold Change
1
2
3
4
Weeks
*
Veh
Cas
Alsrtb
Cas/Alsrtb

**B**

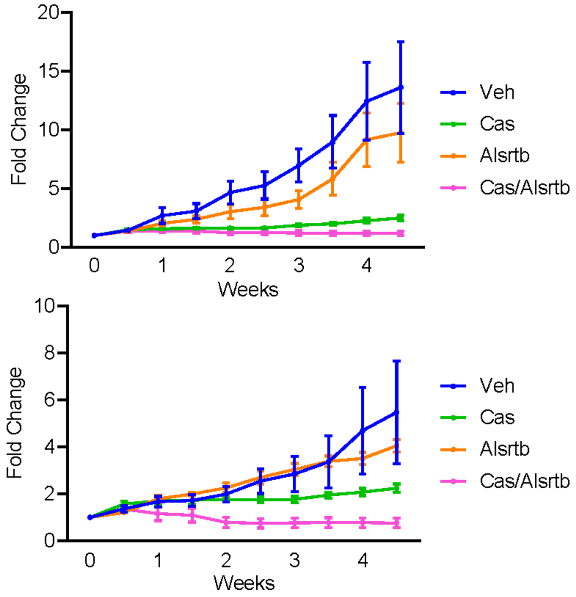
20
15
10
5
0
Fold Change
0
1
2
3
4
Weeks
Veh
Cas
Alsrtb
Cas/Alsrtb
10
8
6
4
2
0
Fold Change
0
1
2
3
4
Weeks
Veh
Cas
Alsrtb
Cas/Alsrtb

**Figure S11. Related to Figure 6. The stem-like/progenitor subpopulation is vulnerable to AURKA inhibition.** Relative change in tumor volume for (A) PC35-1 and (B) VCaP-D (top) and VCaP-16 (bottom) xenografts during four weeks of the indicated treatments. Mice were dosed twice-daily with 20mg/kg of alisertib (Alisertb) or vehicle (Veh). Tumor volume was calculated as an average of the replicates. The change in volume was calculated relative to the "0" time-point (~150mm$^3$ tumor volume). Vehicle n = 5 mice; castration (Cas) n = 5 mice; alisertib n = 5 mice; castration + alisertib n = 5 mice. Error bars, ± SEM. P-values were calculated using the student's t-test, two-tailed, unpaired.

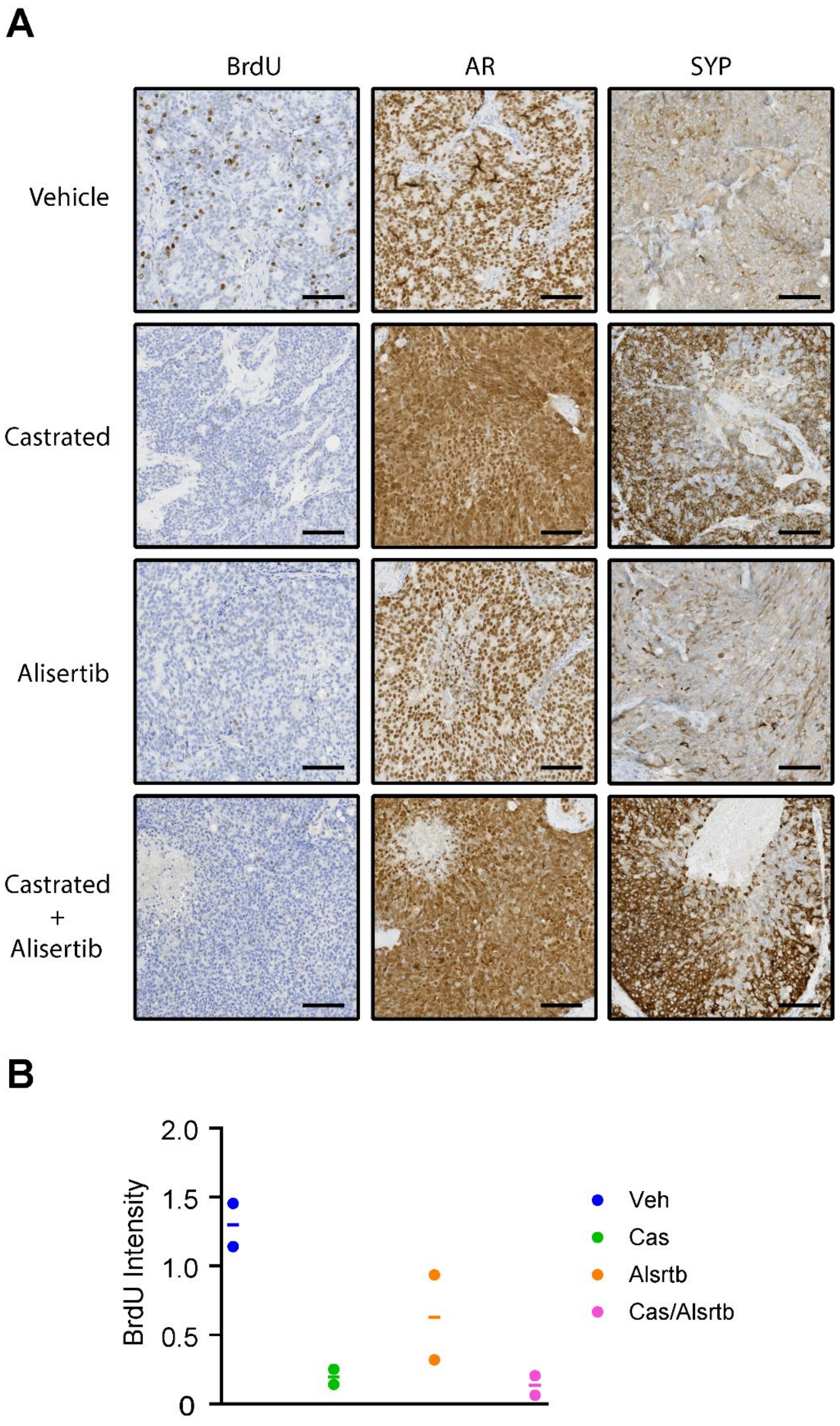
A
BrdU
AR
SYP
Vehicle
Castrated
Alisertib
Castrated
+
Alisertib
B
2.0
1.5
1.0
0.5
0
BrdU Intensity
Veh
Cas
Alsrtb
Cas/Alsrtb

**Figure S12. Organoid-derived xenograft (ODX) tumors respond to castration and treatment with the AURKA inhibitor, alisertib. Related to Figure 6.** (A) Serial sections of PC35-1 ODX tumor tissue from treated (9 weeks) or control mice were stained with antibodies against BrdU, AR or SYP. Mice were injected with BrdU five hours prior to harvest. Scale bars, 100 μM. (B) Two stained sections from each treatment cohort were quantified for BrdU-incorporation. A minimum of 200,000 cells/section were quantified. The average intensity for cells classified as strong Positive (see methods) are plotted. Horizontal line indicates the mean.
